# Evaluations of retrospective frequency and phase correction methods for single-voxel MR spectroscopy at 7T

**DOI:** 10.64898/2026.09.21.753210

**Authors:** Chu-Yu Lee, Jia Xu, Baolian Yang, Ralph Noeske, Vincent A Magnotta

## Abstract

Subject motion and gradient heating-induced frequency and phase offsets result in spectral misalignment during single-voxel MR spectroscopy (SVS) acquisitions. Several methods have been presented to align the spectra, but their performance on a 7T system is unclear. This study aimed to evaluate the practical importance of four retrospective correction methods, namely creatine fitting, residual water, spectral registration, and cross-correlation. SVS data were collected from 127 participants (88/39 female/male, 39±15 years) using a semi-localization by adiabatic selective refocusing sequence at 7T. Changes in the spectral linewidth, signal-to-noise ratio (SNR), similarity (mean similarity matrix (SI_mean_)), and metabolite quantification (concentration, relative Cramer-Rao lower bounds (rCRLB)) after correction were evaluated. A Wilcoxon signed rank test was used to evaluate these changes. The *p*-values were adjusted using the false discovery rate (*q*<0.05) for multiple comparisons. For significant changes, the effect size was further calculated using the Rosenthal formula. The results demonstrated that all correction methods resulted in a significant decrease in the spectral linewidth and significant increases in the spectral SNR and SI_mean_. The mean changes by using the four methods were 0.76-0.99 Hz for the spectral linewidth change (effect sizes:0.73-0.86), 5.79-6.95 for the SNR change (effect size:0.60-0.72), and 0.03-0.03 for the SI_mean_ change (effect size:0.87-0.87). The correction also significantly increased the estimates of the metabolite concentrations of PCr, Gln, Glu, GPC, Ins, NAA, Scyllo, and Tau (effect size:0.25-0.57), and significantly decreased the rCRLBs (effect size:0.26-0.42) of Scyllo, Tau, and Gly. These results indicate that each method provides practical benefits in improving the spectral linewidth, SNR, and similarity by showing a strong to very strong effect of the improvements (effect size>0.60). The correction only showed a weak to moderate effect of the changes in the metabolite quantification (effect size<0.60), suggesting that whether the correction results in more reliable metabolite quantification requires further validation.

## Introduction

Proton MR spectroscopy allows non-invasive quantification of neurochemicals in the brain[1–4]. It has been shown to be useful in characterizing neurometabolic changes in normal aging [1], neurodegenerative diseases [2], brain tumors [3], and psychiatric disorders [4]. For more accurate metabolite quantification, single-voxel MR spectroscopy (SVS) requires multiple averages to boost the signal-to-noise ratio (SNR), which relies on the spectra of the transients being aligned [5]. However, the frequency and phase of each transient changes over time due to gradient heating [6] and subject motion [7]. These frequency and phase offsets lead to misaligned spectra, and direct averaging results in decreased SNR, broader linewidth of the spectrum, and increased biases and uncertainty in metabolite quantification. It is thus important to correct the frequency and phase offsets of each transient before averaging [5].

The frequency and phase offsets can be corrected using retrospective methods without pulse sequence modifications [7–17]. In the retrospective methods, the frequency and phase offsets of each transient are measured using a creatine peak (Cre) [11], residual water peak (RW) [8], or an extracted spectrum, such as in spectral registration (SR) [12] and cross-correlation (Xcorr) [14,15] methods. The measured frequency and phase offsets are then removed from each transient before averaging. A similar concept has been adapted to a deep learning framework [18–22]. Those retrospective methods are typically evaluated based on their ability to account for known frequency and phase variations. This is achieved by adding frequency and phase offsets to in vivo or simulated spectra, followed by comparing the measured offsets to the true offsets to quantify the accuracy of the correction. Adding frequency and phase offsets to the spectra is a good approximation of the gradient heating-induced effects that have minimal impact on the spectral linewidth and line shape of each transient [6]. However, it fails to account for subject motion-induced effects, such as line broadening or line-shape distortion of each transient [7,8,23]. Furthermore, unstable residual water signals can arise from gradient heating or subject motion-induced frequency offsets [23,24]. Therefore, it is important to evaluate the real-world performance of these correction methods using in vivo data.

For in vivo data with unknown frequency and phase offsets, the retrospective correction methods are evaluated based on the changes in the spectral linewidth, SNR, similarity and metabolite quantification after correction [12,15,22,25]. Several studies have used a large dataset (N>50) to evaluate the performance of different frequency and phase correction methods on 3T systems. One study used SVS data from 53 human subjects to show decreases in the spectral linewidth and changes in the metabolite concentrations after correction using the Cre, RW, and SR methods [25]. Another study used 73 human SVS datasets from the Big GABA dataset [26] to compare the model-based correction and SR methods under different SNR levels [16]. On 7T systems, several human studies have applied the Cre, SR, and Xcorr methods to correct frequency and phase offsets for SVS data [27–30]. However, the real-world performance of the different correction methods is unclear. The performance of the correction may be influenced by the broader spectral linewidth at 7T [31].

Thus the aim of this study was to evaluate the practical importance of four retrospective correction methods, namely Cre [11], RW [8], SR [12], and Xcorr [14,15] by using SVS data from more than 100 participants on a 7T system, based on changes in spectral linewidth, SNR, similarity, and metabolite quantification after correction. An additional aim was to simulate SVS spectra over the ranges of the spectral SNR and linewidth at 7T to evaluate the accuracy of the frequency and phase measurements.

## Materials and Methods

### Simulated data

Simulations were performed using the FID-A software [32] to evaluate the accuracy of the frequency and phase measurements of the correction methods. An SVS experiment was simulated with parameters identical to those used in an in vivo semi-localization by adiabatic selective refocusing (semi-LASER) experiment [33] (TE=30 ms, spectral width=5000 Hz, 2048 data points, 32 transients, B_0_=7T, GOIA-WURST refocusing pulses) to generate the spectra of 20 metabolites: ascorbate (Asc), alanine (Ala), aspartate (Asp), creatine (Cr), phosphocreatine (PCr), γ-aminobutyric acid (GABA), glucose (Glc), glutamine (Gln), glutamate (Glu), glycerophosphorylcholine (GPC), phosphorylcholine (PCh), glutathione (GSH), lactate (Lac), myo-inositol (Ins), N-acetylaspartate (NAA), N-acetylaspartylglutamate (NAAG), scyllo-inositol (Scyllo), taurine (Tau), glycine (Gly), and phosphorylethanolamine (PE). The signals of the residual water and macromolecules were also simulated by modeling the signals as singlets at water frequency and macromolecule resonances [34]. The concentrations of the metabolites, residual water, and macromolecules were set to be the median concentrations of those in the in vivo measurements (Supplementary Table 1). A line broadening factor (13.27 Hz) was applied to the simulated spectrum to generate a 16-Hz NAA linewidth that was close to the median NAA linewidth of the in vivo measurements. Additionally, normally distributed random noise was added to the simulated spectrum to generate an SNR of the NAA peak (NAA SNR=128) that was close to the median NAA SNR of the in vivo measurements. The NAA SNR was defined as the ratio of the NAA peak amplitude to the standard deviation (SD) of the noise between -3.5 and 0 ppm. The NAA linewidth and NAA SNR were referred to as the spectral linewidth and SNR, respectively, herein.

Thirty-two noisy transients were simulated to represent a single dataset. Frequency and phase offsets were added to each simulated transient. The frequency offsets were specified as a linear offset across the 32 transients, starting from 0 Hz to a maximum randomly selected between 0 and 10 Hz. Normally distributed noise (0±1 Hz) was added to the linear frequency offset. The phase offsets were uniformly distributed over an interval [0 20]°. One thousand datasets containing these random frequency and phase offsets were simulated.

To study the effects of the spectral SNR and linewidth on the correction, the SD of the noise and line broadening factor were increased, respectively, in two separate experiments. In one experiment, the SD of the noise was increased up to 2.3 times, corresponding to the SNR decreasing from 128 to 55.65 with the line broadening factor kept the same. In the other experiment, the line broadening factor was increased from 13.27 Hz to 22.27 Hz, corresponding to the spectral linewidth increasing from 16 to 25 Hz, with the SD of the noise kept the same. For each condition of the spectral SNR and linewidth, the simulation was repeated 20 times.

### Phantom data

Phantom SVS data were collected to measure the extent of the gradient heating-induced frequency offsets on the 7T system after a one-hour fMRI scan. Non-water-suppressed transients were collected from a spherical MRS Braino phantom (GE Healthcare, Milwaukee, WI, USA) after a one-hour fMRI scan using the same sequence as in the in vivo experiment. The data collection was repeated in the morning but without prior MRI scans for 12 hours as a baseline. The acquisition parameters were as follows: TE=30 ms, TR=4000 ms, spectral width=5000 Hz, 2048 data points, and 128 non-water-suppressed transients. For each condition (with heating or baseline), the experiment was repeated five times.

### In vivo data

The SVS data were collected from 144 participants (100/44 female/male; 39±15 years old) who were enrolled in a study of bipolar disorder from January 2023 to July 2024. This study was approved by the Institutional Review Board. Informed consent was obtained to participate in the study that included MRI and SVS measurements from all participants. The participants were scanned on a GE SIGNA 7T system (GE Healthcare, Milwaukee, WI, USA) using a NOVA 2-channel transmit/32-channel receive coil. The 20×20×20 mm^3^ volume-of-interest (VOI) was placed on the anterior cingulate cortex using the semi-LASER localization combined with VAPOR water suppression and 3D outer volume suppression [33]. A vendor-supplied high-order B_0_ shimming routine was performed on the VOI before the SVS acquisition. The acquisition parameters were as follows: TE=30 ms, TR=5000-6430 ms, spectral width=5000 Hz, 2048 data points, 32 transients, and 2 non-water-suppressed transients.

### Frequency and phase correction

For the in vivo data, following coil combination [35], each of the four frequency and phase correction methods implemented in Matlab (The Mathworks, Natick, Massachusetts, USA) was applied to measure the frequency and phase offsets of the transients, as briefly described below:

1. Cre method [11,12]: the signals of each transient were zero-filled to 20480 points and apodized with a 2-Hz exponential filter. The spectrum of the transient was windowed from 2.72 to 3.12 ppm to extract the creatine peak. The peak was fitted with a Lorentzian function to determine the frequency and phase offsets.
2. RW method [8]: the signals of each transient were zero-filled to 20480 points and apodized with a 2-Hz exponential filter. The spectrum of the transient was windowed from 4.4 to 5 ppm to extract the residual water peak. The peak location determined the frequency offset. After the removal of the frequency offset, the remaining phase of the first data point of the transient determined the phase offset.
3. SR method [12]: the signals with a time-domain SNR above 5 (by default in the FID-A software [32]) and a frequency range of 1.8 to 3.6 ppm were extracted from each transient. The extracted signals were then aligned to those of a reference transient by adjusting the frequency and phase offsets using the nonlinear fitting in Matlab. The frequency and phase offsets that resulted in the best alignment were the measured frequency and phase offsets for the transient.
4. Xcorr method [15]: the signals of each transient were zero-filled to 20480 points and apodized with a 2-Hz exponential filter. The spectrum of each transient was windowed from 1.8 to 3.6 ppm and cross-correlated with that of a reference transient. The shift and phase of the cross-correlation peak determined the measured frequency and phase offsets, respectively, for each transient. The reference transient of each dataset was defined as the transient that was closest to the median of the transients.

For each correction method, the measured frequency and phase offsets were removed from each of the transients to generate the frequency and phase corrected transients.

For the phantom data with non-water-suppressed transients, the RW method was used to measure the gradient heating-induced frequency offsets.

### Metabolite quantification

Metabolite quantification was performed for the in vivo data using LCModel [36]. The basis consisting of 20 metabolites was simulated using the FID-A software [32] as described in the simulation. The linewidth of the singlet in the basis set was 3.87 Hz (FWHMBA=0.013 ppm). The LCModel fit to the spectrum was performed between 0.3 and 4.1 ppm to obtain estimates of metabolite concentrations and relative Camér–Rao lower bounds (rCRLB). The non-water-suppressed transient was used for water scaling and eddy current correction. The water relaxation attenuation (ATTH2O) was set to 0.56 according to a previously reported T_2_ relaxation time of water at 7T [37]. WCONC was set to 55556. No corrections for partial volume and relaxation effects were performed for this analysis.

### Quality control

Data with poor shimming (linewidth of non-water-suppressed transient>25 Hz) [31], poor water suppression (ratio of water to NAA peaks>5), broad spectral linewidth (spectral linewidth>30 Hz) or low SNR (SNR<28) were excluded from the analysis [38]. Motion-corrupted transients were defined as the transients that showed line broadening or line-shape distortion resulting from subject motion which cannot be recovered by the frequency and phase correction. They were identified by computing the correlation coefficient (Corr) between the spectra of the i_th_ transient (S_i_(f)) and a reference transient (S_ref_(f)) over a range from 1.8 to 3.6 ppm [22]:

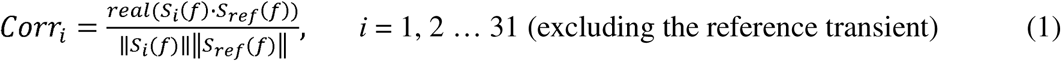

Transients with a Corr less than 0.8 after correction were considered as motion-corrupted transients and were removed from the dataset.

### Evaluation

The accuracy of the frequency and phase measurements for the simulation data with known frequency and phase offsets was evaluated. The accuracy was determined by the frequency and phase errors, defined as the mean absolute differences between the measured and true offsets in each dataset [12,22]. The evaluation also focused on changes in the spectral SNR and linewidth of the averaged spectrum, as well as the changes in the mean of the similarity matrix (SI_mean_) after correction. The SI_mean_ computes the spectral similarity of the N transients in each dataset [22]:

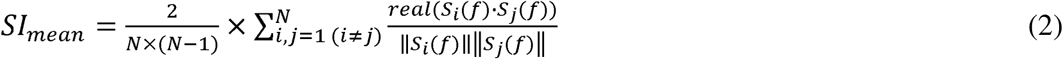

Finally, changes in the estimates of the metabolite concentrations and rCRLB after correction were evaluated.

For the Cre and RW methods that utilized the single spectral peak information, the stability of the peak amplitude within a dataset was investigated by computing the within-dataset SD of the peak amplitude before correction.

### Statistical Analysis

The changes in the spectral SNR, linewidth, and SI_mean_ after correction were evaluated using a Wilcoxon signed rank test. For the in vivo data, the changes in the estimates of metabolite concentrations and rCRLB after correction were also evaluated using a Wilcoxon signed rank test. The Wilcoxon sign-rank test was used to evaluate the changes after correction without making assumptions about the data distribution. The normality of the data distribution was evaluated using a one-sample Kolmogorov-Smirnov test.

Within each dataset, only the estimates of the metabolite concentrations with a rCRLB<50% before and after correction were included in the comparison. Only the comparisons of the metabolite concentrations reaching 50 or more pairs out of the 127 datasets were included in the statistical analysis to ensure sufficient statistical power. All *p*-values were adjusted using the false discovery rate (FDR) [39] with a significance level (*q*<0.05) to account for multiple comparisons (David Groppe (2026) function fdr_bh: https://www.mathworks.com/matlabcentral/fileexchange/27418-fdr_bh, MATLAB Central File Exchange). In total, 760 comparisons were evaluated in this study. The comparisons included 380 correction-induced changes evaluated using the Wilcoxon sign-rank test: 240 changes for the simulation data (20 conditions of spectral quality×4 correction methods×3 (linewidth, SNR, and SI_mean_) and 140 changes for the in vivo data (4 correction methods×3 (linewidth, SNR, and SI_mean_) +16 metabolites×4 correction methods×2 (concentration and rCRLB)). The comparisons also included the data distributions of all the 380 changes evaluated using the one-sample Kolmogorov-Smirnov test.

For significant changes, the effect size was used to quantify the magnitude of the change. The effect size was calculated using the Rosenthal formula, defined as the z statistic divided by the square root of the sample size [40,41]. The effect size was presented as an absolute value and interpreted as [42]: 0.2 or lower, very weak, 0.2 to 0.4, weak, 0.4 to 0.6, moderate, 0.6 to 0.8, strong, 0.8 or higher, very strong. The 95% confidence interval (CI) of the effect size was computed using bootstrap resampling with 10000 iterations.

## Results

For the simulation, the data distributions of the linewidth and SI_mean_ changes significantly deviated from a normal distribution. For the in vivo data, the distribution of the linewidth change significantly deviated from a normal distribution. The distributions of the concentration changes of Cr, PCr, and PE, as well as the rCRLB changes of Asp, Cr, PCr, GABA, Gln, Glu, GPC, GSH, Ins, NAA, Tau, Gly, and PE, significantly deviated from a normal distribution. To ensure that the statistical tests used to evaluate the changes had the same statistical power and that the effect size was computed in an identical manner, the Wilcoxon sign-rank test was used to evaluate the changes throughout this study.

### Simulation data

For the noise-free spectra, all the correction methods had average frequency and phase measurement errors below 0.2 Hz/0.5° (Supplementary Fig. 1). With added noise, the shape of the simulated spectrum agreed with that of the in vivo spectrum (Supplementary Fig. 2). The frequency and phase offsets were further added to the spectra to evaluate each correction method (Fig. 1). Over the conditions of the spectral SNR and linewidth (Fig. 2), the SR method had the most accurate frequency measurement (mean error: 0.27 Hz), followed by the Xcorr (mean error: 0.4 Hz), Cre (mean error: 0.58 Hz), and RW methods (mean error: 0.77 Hz). The Xcorr method had the most accurate phase measurement (mean error: 0.79°), but the accuracy was close to that of the SR (mean error: 0.83°) and RW methods (mean error: 0.86°). The Cre method had the lowest accuracy of the phase measurement (mean error: 3.42°).

**Figure 1:**
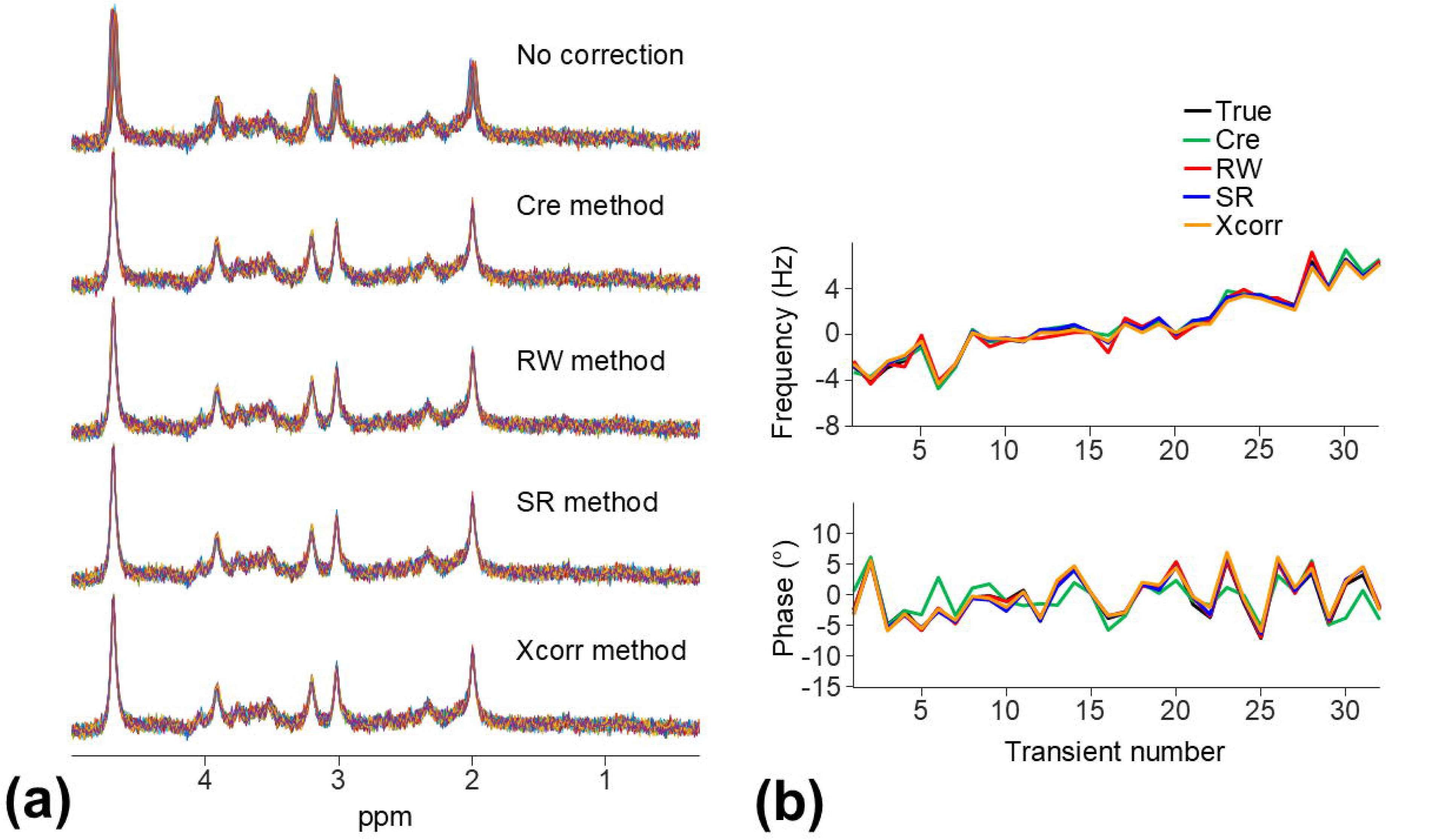
Simulation: **(a)** An example of applying the frequency and phase correction to the 32 simulated noisy spectra (spectral linewidth: 16 Hz and SNR: 126.54) with known frequency and phase offsets. **(b)** The ranges of the simulated frequency and phase offsets within this dataset were 10.85 Hz/12.7°. The average frequency and phase errors using each of the four correction methods were 0.27 Hz/1.78° (Cre), 0.30 Hz/0.38° (RW), 0.13 Hz/0.51° (SR), and 0.24 Hz/0.62° (Xcorr).

**Figure 2:**
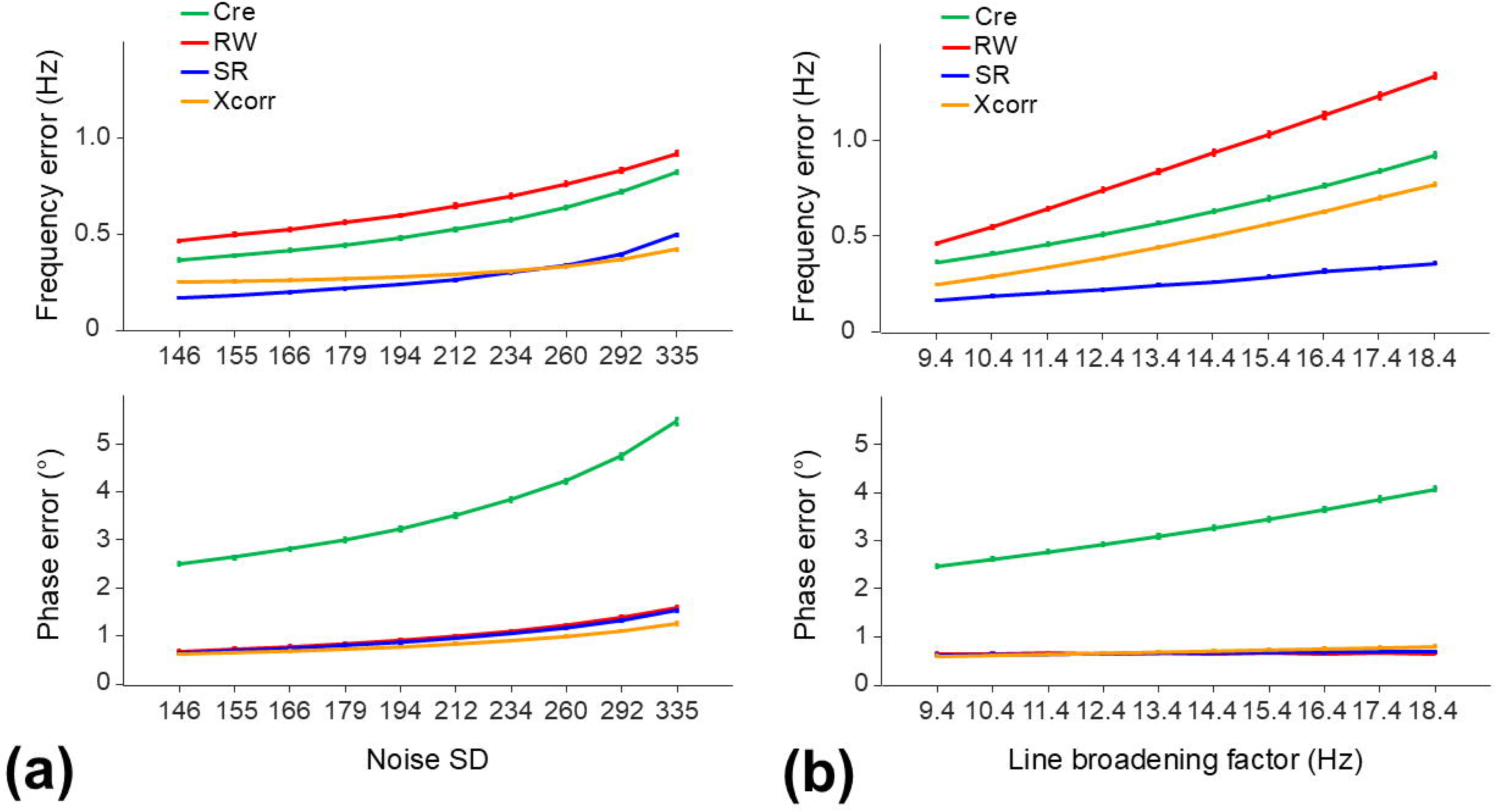
Simulation: The frequency and phase measurement errors (mean ± SD over the 20 repeated experiments) using each of the four correction methods under the specified conditions of the spectral SNR (**a**) and linewidth (**b**). The increased noise SD from 146 to 335 in **a** corresponds to a decrease in the spectral SNR from 128 to 55.65 with the line broadening factor (9.4 Hz) kept the same. The increased line broadening factor from 13.27 Hz to 22.27 Hz in **b** corresponds to an increase in the spectral linewidth from 16 to 25 Hz with the noise SD (146) kept the same.

The correction using each method resulted in significant decreases in the spectral linewidth, and significant increases in the spectral SNR and SI_mean_ under each condition of the spectral SNR and linewidth (Supplementary Fig. 3). The effect sizes of these changes were smaller when the noise SD and line broadening factor were larger (Fig. 3). The correction using each of the methods showed a strong to very strong effect of the spectral linewidth, SNR, and SI_mean_ changes, except for the spectral SNR change when the line broadening factor was large. When the line broadening factor was 13.4 Hz or larger (spectral linewidth of 20 Hz), the correction showed a weak to moderate effect of the spectral SNR change. The effect size and 95% CI under each condition of the spectral SNR and linewidth are reported in Supplementary Tables 2 and 3.

**Figure 3:**
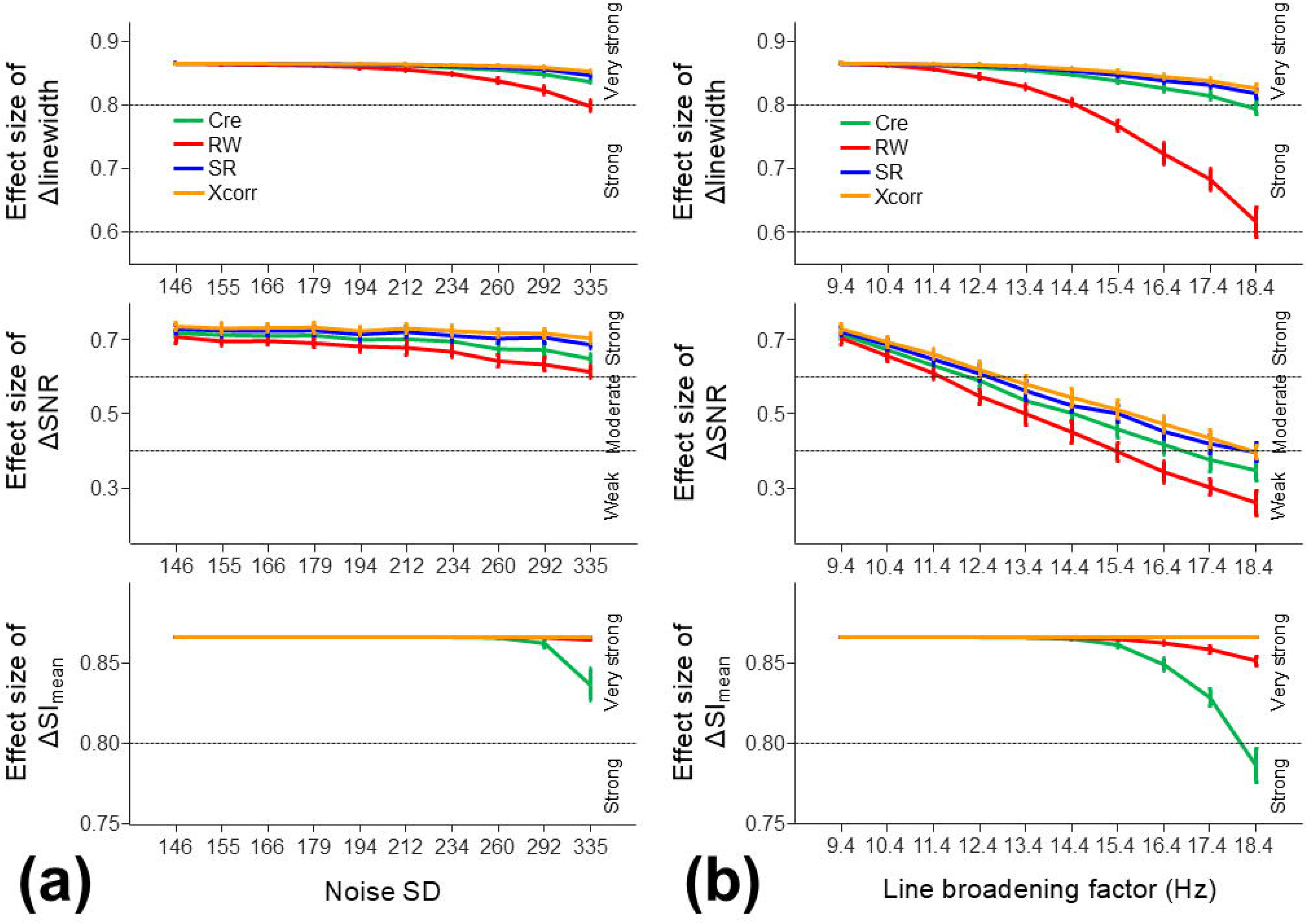
Simulation: The effect size (mean ± SD over the 20 repeated experiments) of the spectral linewidth, SNR, and SI_mean_ changes following the frequency and phase correction using each of the four methods under the specified conditions of the spectral SNR (**a**) and linewidth (**b**). The increased noise SD from 146 to 335 in **a** corresponds to a decrease in the spectral SNR from 128 to 55.65 with the line broadening factor (9.4 Hz) kept the same. The increased line broadening factor from 13.27 Hz to 22.27 Hz in **b** corresponds to an increase in the spectral linewidth from 16 to 25 Hz with the noise SD (146) kept the same. The effect size was computed using the Rosenthal formula for the Wilcoxon signed rank test. The effect size was interpreted as [42]: 0.2 or lower, very weak, 0.2 to 0.4, weak, 0.4 to 0.6, moderate, 0.6 to 0.8, strong, 0.8 or higher, very strong.

### Phantom data

The measured rate of the gradient heating-induced frequency offset was 0.11±0.05 Hz/min after a one-hour fMRI scan (Supplementary Fig. 4). The offset rate was a small increase from the baseline (0.06±0.05 Hz/min) on the 7T system equipped with a superconducting shim coil. For the duration of an in vivo SVS acquisition (2 min 48 s), the gradient heating-induced frequency offset was estimated to be 0.32±0.14 Hz.

### In vivo data

According to the exclusion criteria, 17 datasets were excluded from the analysis: linewidth of non-water-suppressed transient>25 Hz (n=12), ratio of water to NAA peaks>5 (n=3), spectral linewidth>30 Hz (n=12) or SNR<28 (n=1). The remaining 127 datasets (88/39 female/male; 39±15 years old) were used to evaluate each correction method. The median spectral linewidth was 15.98 Hz; range: 10.32-29.76 Hz. The median spectral SNR was 142.52; range: 62.2-237.24.

Spurious echoes were identified between 4.1-4.5 ppm or 4.9-5.3 ppm in most of the datasets (supplementary Fig. 5). These frequency ranges were largely outside the ranges used for the frequency and phase correction, e.g., 2.72-3.12 ppm (Cre), 4.4-5 ppm (RW), and 1.8-3.6 ppm (SR and Xcorr), as well as 0.3-4.1 ppm for the metabolite quantification.

Among the 127 datasets, only one transient of a dataset was identified as a motion-corrupted transient (Corr<0.8 after correction) and was removed (Fig. 4). Most of the datasets showed increases in the Corr after correction: 17% of the datasets originally contained transients with a Corr less than 0.8 and this was reduced to less than 1% after correction regardless of the correction method.

**Figure 4:**
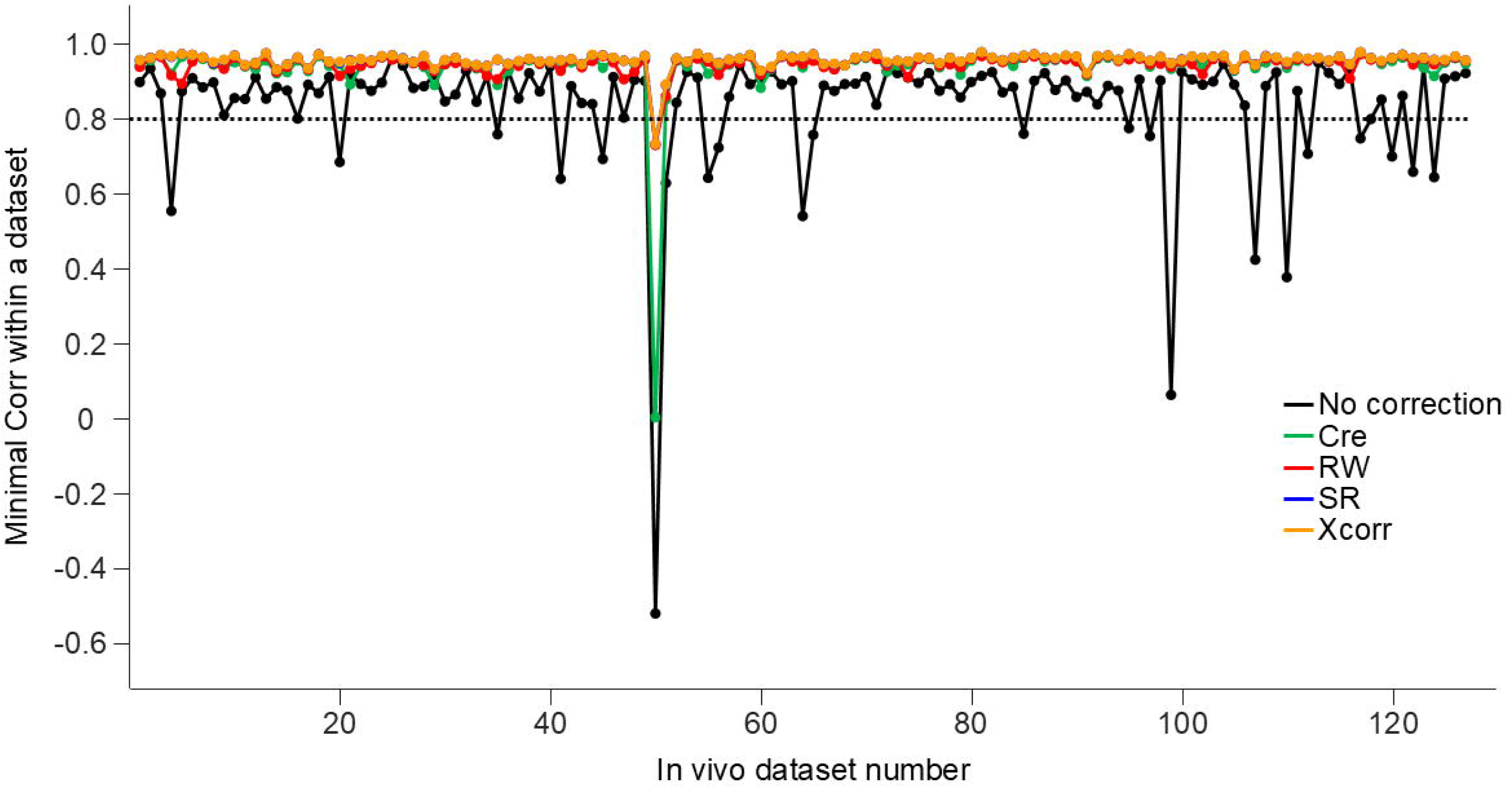
In vivo data: The minimal correlation coefficient (Corr) within each of the 127 in vivo datasets without and with the frequency and phase correction using each of the four methods. The Corr measures the correlation between the spectra of the transient and a reference transient within a dataset. The dash line indicates the threshold to determine the motion-corrupted transients (Corr < 0.8 after correction).

Over the 127 datasets, the average ranges of the measured frequency and phase offsets within a dataset were 6.76 Hz/30.68° (Cre), 6.89 Hz/35.45° (RW), 6.07 Hz/35.73° (SR), and 5.54 Hz/35.68° (Xcorr). The ranges of the average frequency and phase offsets per transient were 0.46-4.61 Hz/2.60-15.29° (Cre), 0.49-5.14 Hz/2.43-13.84° (RW), 0.30-3.67 Hz/2.43-13.99° (SR), and 0.02-3.49 Hz/2.30-13.79° (Xcorr).

Fig. 5 illustrates effective alignment of the spectra after correction in a representative dataset. The correction resulted in a significant decrease in the spectral linewidth and significant increases in the spectral SNR and SI_mean_ (Fig. 6). The mean changes of the spectral linewidth were 0.99 (Cre), 0.76 (RW), 0.96 (SR), and 0.95 Hz (Xcorr) with a strong to very strong effect size; effect sizes: 0.86, 95% CI [0.85, 0.87] (Cre), 0.73, 95% CI [0.64, 0.80] (RW), 0.84, 95% CI [0.80, 0.86] (SR), and 0.84, 95% CI [0.82, 0.86] (Xcorr). The mean changes of the SNR were 6.38 (Cre), 5.79 (RW), 6.91 (SR), and 6.95 (Xcorr) with a strong effect size; effect sizes: 0.71, 95% CI [0.61, 0.79] (Cre), 0.60, 95% CI [0.47, 0.70] (RW), 0.70, 95% CI [0.61, 0.78] (SR), and 0.72, 95% CI [0.63, 0.79] (Xcorr). The mean changes of the SI_mean_ were 0.03 (Cre), 0.03 (RW), 0.03 (SR), and 0.03 (Xcorr) with a very strong effect size; effect sizes: 0.87, 95% CI [0.86, 0.87] (Cre), 0.87, 95% CI [0.87, 0.87] (RW), 0.87, 95% CI [0.87, 0.87] (SR), and 0.87, 95% CI [0.87, 0.87] (Xcorr).

**Figure 5:**
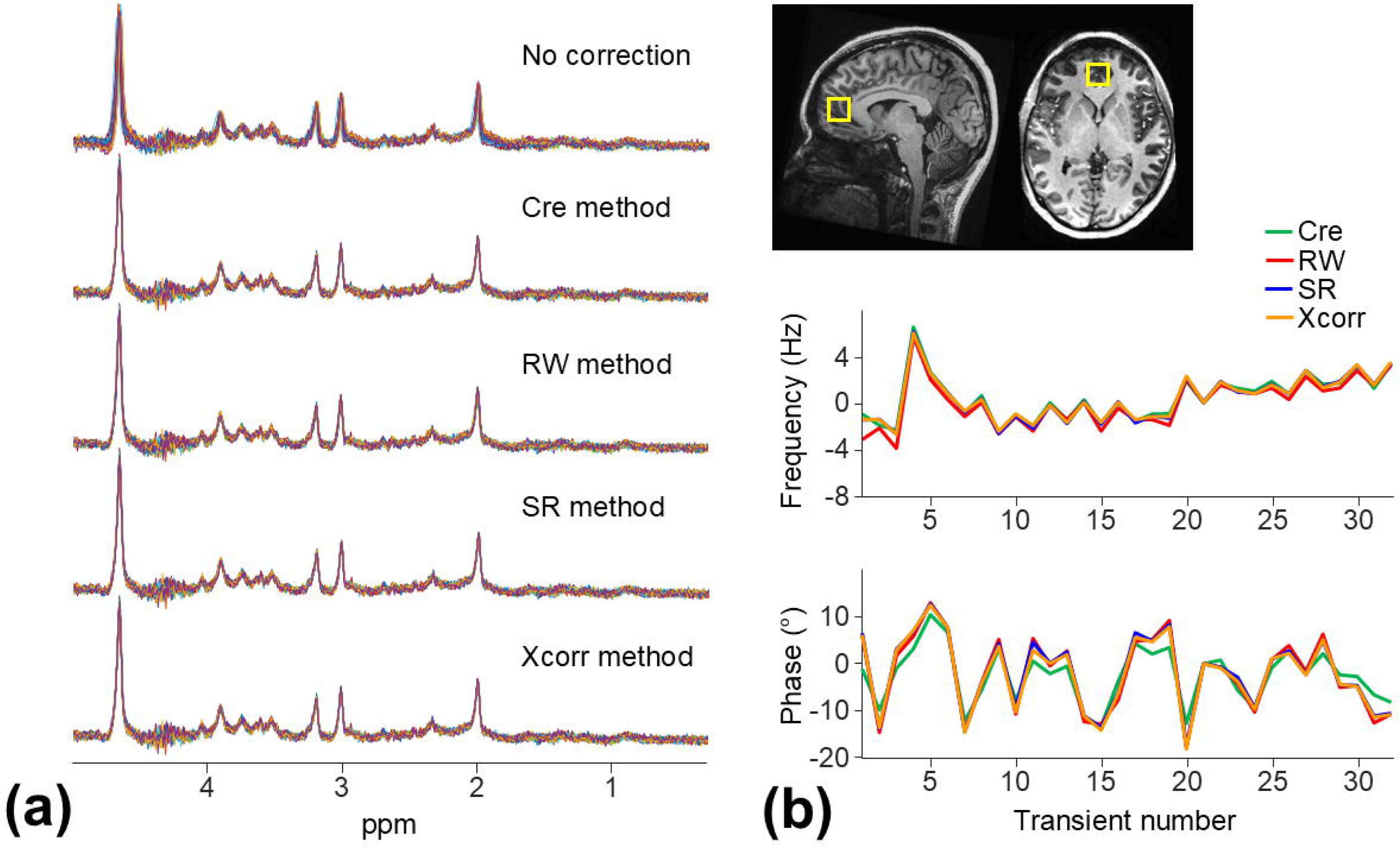
In vivo data: An example of applying the frequency and phase correction to 32 spectra of an in vivo dataset (spectral linewidth: 12.15 Hz and SNR: 184.36) using each of the correction methods. The ranges of the measured frequency and phase offsets within this dataset were 9 Hz/23.69° (Cre), 9.52 Hz/30.7° (RW), 8.76 Hz/30.27° (SR), and 8.54 Hz/30.5° (Xcorr).

**Figure 6:**
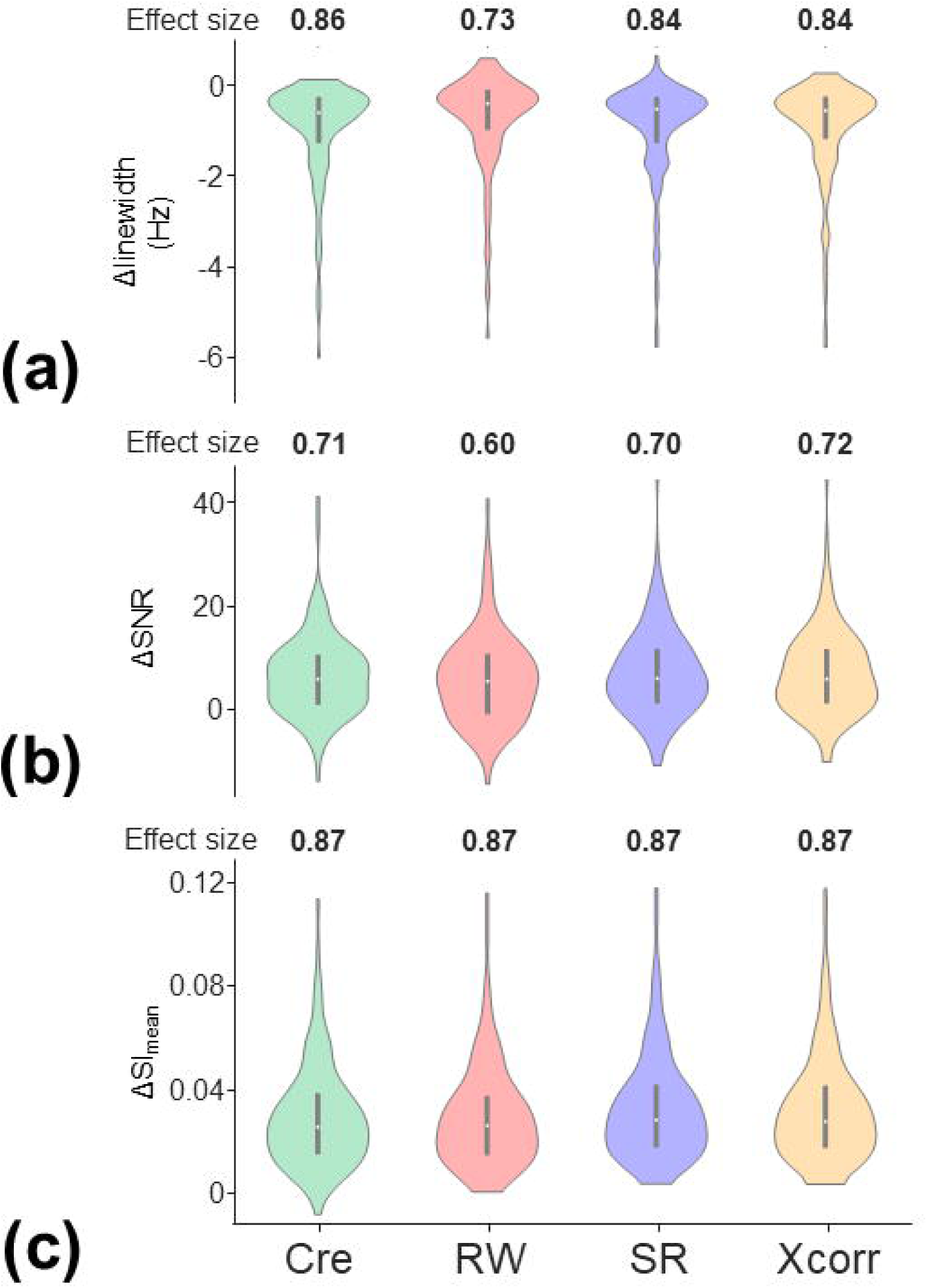
In vivo data: Distributions and effect sizes of the changes of the spectral linewidth (a), SNR (b), and SI_mean_ (c) of the 127 in vivo datasets after the frequency and phase correction using each of the four methods. The gray line indicates the 25th percentile, median, and 75th percentile of the changes. The effect size was computed using the Rosenthal formula.

Over the 127 datasets, the average within-dataset SD of the residual water peak amplitude was 2.15 times higher than that of the creatine peak amplitude (Fig. 7).

**Figure 7:**
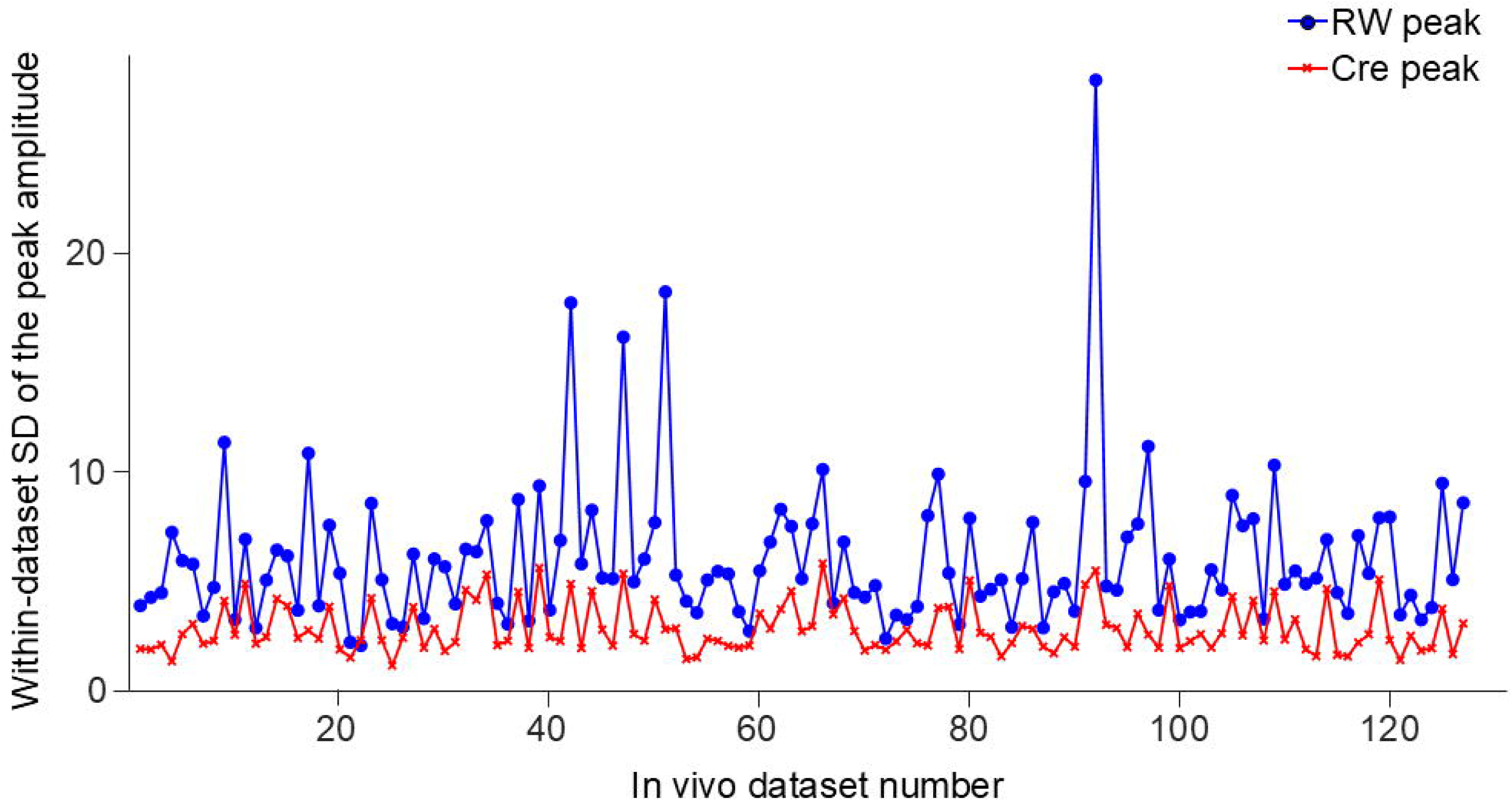
In vivo data: The standard deviation (SD) of the Cre and RW peak amplitude within each of the 127 in vivo datasets before the frequency and phase correction. Over the 127 datasets, the average SD of the Cre peak amplitude within a dataset was 2.86; range: 1.17-5.85. The average SD of the RW peak amplitude within a dataset was 6.03; range: 2.08-28.

Fig. 8 illustrates an example of the LCModel fit to an uncorrected in vivo spectrum for the metabolite quantification. Tables 1 and 2 list the metabolites showing significant differences in the metabolite quantification after correction. The correction resulted in significant increases in the measured concentrations of PCr, Gln, Glu, GPC (except for the RW method), Ins, NAA, Scyllo, and Tau (Table 1) with a weak to moderate effect size; effect sizes: 0.25-0.57. The corrections also resulted in significant decreases in the rCRLB of Scyllo, Tau (except for the Cre, RW, and SR methods), and Gly (except for the RW and SR methods) (Table 2) with a weak to moderate effect size; effect sizes: 0.26-0.42. The effect size and its 95% CI of the changes in the measured metabolite concentration and rCRLB were reported in Tables 1 and 2.

**Figure 8:**
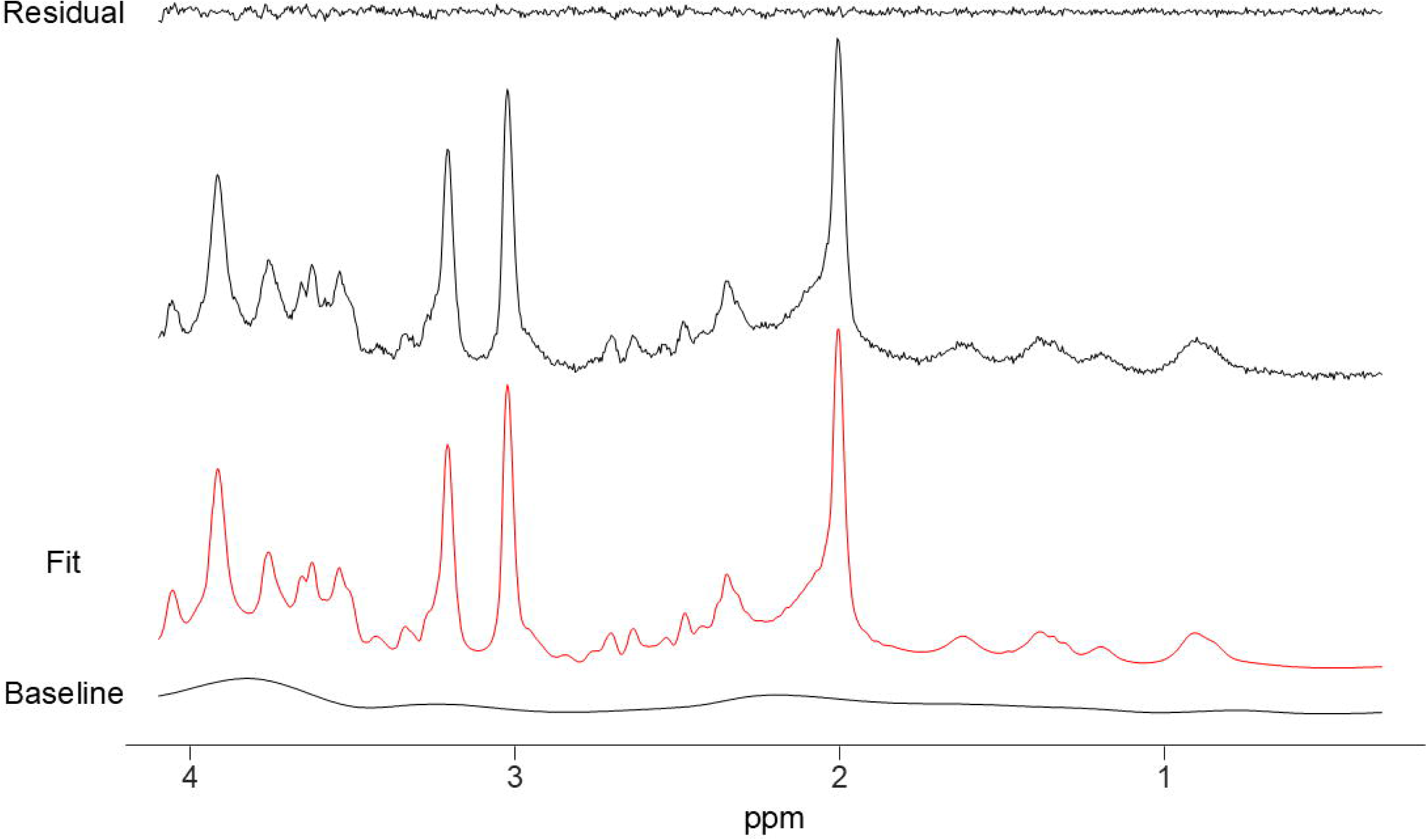
In vivo data: An example of the LCModel fit to an uncorrected in vivo spectrum (averaged over 32 transients). The fit is shown along with the residuals and baseline.

**Table 1:** In vivo data: List of metabolites with estimates of concentrations (in mM) showing significant changes following the frequency and phase correction using each of the four methods. The estimates of the metabolite concentrations were obtained through the LCModel fit to the in vivo spectra. N indicates the number of comparisons which were evaluated using a Wilcoxon signed rank test. The *p*-values were adjusted using the false discovery rate with a significance level (*q* < 0.05). The effect size was calculated using the Rosenthal formula. The 95% CI of the effect size was computed using bootstrap resampling with 10000 iterations.

| Metabolite | PCr | Gln | Glu | GPC | Ins | NAA | Scyllo | Tau |
| --- | --- | --- | --- | --- | --- | --- | --- | --- |
| N | 127 | 126 | 127 | 125 | 127 | 127 | 79 | 126 |
| Conc (no corr) | 5.27 | 1.65 | 9.75 | 1.39 | 6.82 | 10.92 | 0.22 | 2.05 |
| Conc (Cre) | 5.43 | 1.68 | 9.82 | 1.44 | 6.88 | 11.01 | 0.23 | 2.09 |
| *q*-value | **<0.00001** | **0.004** | **<0.00001** | **0.03** | **0.00009** | **<0.00001** | **0.008** | **<0.00001** |
| Effect size | **0.49** | **0.32** | **0.50** | **0.26** | **0.40** | **0.48** | **0.38** | **0.45** |
| 95% CI | **[0.35, 0.62]** | **[0.15, 0.48]** | **[0.35, 0.64]** | **[0.09, 0.42]** | **[0.24, 0.55]** | **[0.33, 0.62]** | **[0.17, 0.56]** | **[0.29, 0.59]** |
| N | 127 | 126 | 127 | 125 | 127 | 127 | 80 | 126 |
| Conc (no corr) | 5.27 | 1.65 | 9.75 | 1.39 | 6.82 | 10.92 | 0.22 | 2.05 |
| Conc (RW) | 5.38 | 1.69 | 9.83 | 1.42 | 6.89 | 11.02 | 0.23 | 2.09 |
| *q*-value | **0.003** | **0.0005** | **<0.00001** | 0.11 | **0.00002** | **<0.00001** | **0.03** | **0.00001** |
| Effect size | **0.32** | **0.37** | **0.55** | 0.23 | **0.42** | **0.51** | **0.34** | **0.44** |
| 95% CI | **[0.16, 0.48]** | **[0.21, 0.52]** | **[0.42, 0.67]** | [0.06, 0.39] | **[0.27, 0.56]** | **[0.36, 0.64]** | **[0.13, 0.52]** | **[0.28, 0.58]** |
| N | 127 | 126 | 127 | 125 | 127 | 127 | 79 | 126 |
| Conc (no corr) | 5.27 | 1.65 | 9.75 | 1.39 | 6.82 | 10.92 | 0.22 | 2.05 |
| Conc (SR) | 5.42 | 1.68 | 9.83 | 1.42 | 6.89 | 11.02 | 0.23 | 2.09 |
| *q*-value | **<0.00001** | **0.008** | **<0.00001** | **0.04** | **0.00001** | **<0.00001** | **0.005** | **0.00009** |
| Effect size | **0.45** | **0.30** | **0.50** | **0.25** | **0.44** | **0.56** | **0.40** | **0.40** |
| 95% CI | **[0.30, 0.59]** | **[0.13, 0.46]** | **[0.35, 0.63]** | **[0.08, 0.42]** | **[0.29, 0.58]** | **[0.42, 0.67]** | **[0.20, 0.58]** | **[0.24, 0.55]** |
| N | 127 | 126 | 127 | 125 | 127 | 127 | 79 | 126 |
| Conc (no corr) | 5.27 | 1.65 | 9.75 | 1.39 | 6.82 | 10.92 | 0.22 | 2.05 |
| Conc (Xcorr) | 5.43 | 1.68 | 9.84 | 1.43 | 6.9 | 11.03 | 0.23 | 2.09 |
| *q*-value | **<0.00001** | **0.0007** | **<0.00001** | **0.03** | **<0.00001** | **<0.00001** | **0.0008** | **<0.00001** |
| Effect size | **0.47** | **0.36** | **0.55** | **0.27** | **0.47** | **0.57** | **0.45** | **0.45** |
| 95% CI | **[0.33, 0.61]** | **[0.20, 0.51]** | **[0.41, 0.67]** | **[0.10, 0.43]** | **[0.32, 0.61]** | **[0.44, 0.69]** | **[0.25, 0.63]** | **[0.30, 0.60]** |

**Table 2:** In vivo data: List of metabolites with relative Camér–Rao lower bounds (rCRLBs) of the concentration estimates (in %) showing significant changes following the frequency and phase correction using each of the four methods. The rCRLBs were obtained through the LCModel fit to the in vivo spectra. N indicates the number of comparisons which were evaluated using a Wilcoxon signed rank test. The *p*-values were adjusted using the FDR with a significance level (*q* < 0.05). The effect size was calculated using the Rosenthal formula. The 95% CI of the effect size was computed using bootstrap resampling with 10000 iterations.

| Metabolite | Scyllo | Tau | Gly |
| --- | --- | --- | --- |
| N | 79 | 126 | 125 |
| rCRLB (no corr) | 24.32 | 10.06 | 13.7 |
| rCRLB (Cre) | 23.08 | 9.71 | 13.3 |
| *q*-value | **0.004** | 0.14 | **0.03** |
| Effect size | **0.40** | 0.22 | **0.27** |
| 95% CI | **[0.21, 0.57]** | [0.06, 0.38] | **[0.11, 0.43]** |
| N | 80 | 126 | 125 |
| rCRLB (no corr) | 24.6 | 10.06 | 13.7 |
| rCRLB (RW) | 23.49 | 9.79 | 13.42 |
| *q*-value | **0.02** | 0.20 | 0.19 |
| Effect size | **0.34** | 0.21 | 0.21 |
| 95% CI | **[0.14, 0.52]** | [0.04, 0.37] | [0.04, 0.38] |
| N | 79 | 126 | 125 |
| rCRLB (no corr) | 24.6 | 10.06 | 13.7 |
| rCRLB (SR) | 23.11 | 9.72 | 13.31 |
| *q*-value | **0.01** | 0.17 | 0.09 |
| Effect size | **0.37** | 0.21 | 0.24 |
| 95% CI | **[0.17, 0.54]** | [0.05, 0.37] | [0.07, 0.39] |
| N | 79 | 126 | 125 |
| rCRLB (no corr) | 24.6 | 10.06 | 13.7 |
| rCRLB (Xcorr) | 23.06 | 9.71 | 13.21 |
| *q*-value | **0.002** | **0.04** | **0.01** |
| Effect size | **0.42** | **0.26** | **0.29** |
| 95% CI | **[0.23, 0.59]** | **[0.10, 0.41]** | **[0.13, 0.44]** |

## Discussion

The measured frequency and phase offsets in the in vivo datasets (average ranges:5.54-6.89 Hz, 30.68-35.73°) were larger than those reported in prior human studies undertaken at 3T without prior gradient-intensive MRI scans (average offsets:1.7-3.41 Hz, 15.3°) [16,25]. The measured offsets are primarily attributed to subject motion since they were approximately 20 times larger than the estimated gradient heating-induced frequency offset using a phantom. Thus, these in vivo datasets allow evaluation of the real-world performance of the correction methods at 7T in the presence of realistic frequency and phase offsets.

The in vivo results demonstrated that the correction using each of the methods resulted in a strong to very strong effect in the improvements of the spectral linewidth, SNR and SI_mean_. The differences in the effect size among the Cre, SR, and Xcorr methods were less than 0.02. The RW method showed lower effect sizes of the improvements in the spectral linewidth and SNR by 0.10-0.13 compared with the other three methods. Given the spectral linewidth and SNR of the in vivo data, these differences in the effect size agreed with the trends shown in the simulation. The lower effect sizes shown by the RW method may, in part, have resulted from the lower accuracy of the frequency measurement observed in the simulation. The smallest improvements may have also resulted from the relatively unstable residual water peak amplitude within a dataset compared with the creatine peak amplitude. Subject motion during the acquisition changes the VOI location and disturbs the B_0_ homogeneity [43]. The consequences include ineffective water suppression and contamination from under-suppressed water signals outside the VOI [23,24], which can contribute to an unstable residual water peak amplitude. Thus, the RW method may be more subject to motion-related effects than the other methods.

The subject motion-related effects on the spectra, including line broadening and line-shape distortion, can be mitigated by using prospective motion correction methods [24,44]. However, they cannot be corrected by using retrospective frequency and phase correction. Therefore, it has been recommended that motion-corrupted transients be identified and removed from each dataset [5]. The current study utilized the correlation measure and identified motion-corrupted transients in less than 1% of the 127 in vivo datasets after correction, while they were present in approximately 15% of transients before correction. These results suggest that following the removal of datasets with poor spectral quality, severe subject motion-induced line broadening and line-shape distortion have only limited impact on the remaining datasets and that a strong correlation (Corr>0.8) with the reference transient can be mostly recovered after correction.

Although the mean changes of the spectral linewidth, SNR, and SI_mean_ were small at a group level, the improvements could be substantial at an individual level based on the skewness of the distributions shown in Fig. 3. The improvements in the spectral linewidth, SNR, and SI_mean_ were up to 5.83 Hz, 43.79, and 0.12, respectively. The larger improvements of the spectral linewidth, SNR, and similarity in these datasets could potentially help more data meet the quality control criteria and increase the statistical power for clinical or research SVS studies. Despite the strong effect of the improvements in the spectral linewidth, SNR, and similarity, the correction only showed a weak to moderate effect on the changes in the estimates of the metabolite concentration and rCRLB. Additionally, the lack of reference estimates, e.g., LCM estimates without the effect of the offsets, also prevents this study from establishing the improvement of the metabolite quantification after correction. Therefore, it requires further validation whether the observed changes in the metabolite quantification provide a practical benefit.

The simulation results indicate that each of the methods had different accuracy of the frequency and phase measurements. Several factors could contribute to the observed differences in the accuracy. First, the methods that utilize the multiple metabolite peak information, such as SR and Xcorr methods, are generally more accurate than the methods that utilize the single peak information, such as RW and Cre methods. This is supported by our simulation and a previous simulation study [12]. Second, for the methods that utilize the single peak information, determining the frequency offset from the fitting to the creatine peak was more accurate than determining it from the residual water peak location. On the other hand, the phase offset estimate from the fitting to the creatine peak may be influenced by the presence of other metabolite, macromolecule, and baseline signals, which may contribute to the lower accuracy of the phase measurement using the Cre method.

The simulation demonstrated that the improvement in the spectral SNR after correction was more subject to a broad spectral linewidth compared with the improvements in the spectral linewidth and SI_mean_. As shown in the simulation, the correction showed a weak to moderate effect of the spectral SNR change when the spectral linewidth was larger than 20 Hz. Similarly, the in vivo results showed a smaller improvement in the spectral SNR after correction compared with the improvements in the spectral linewidth and SI_mean_. These results suggest that in the presence of a broad spectral linewidth, the benefit of the correction becomes less notable in the spectral SNR change compared with those shown in the spectral linewidth and SI_mean_ changes.

The ranges of the simulated frequency and phase offsets were determined based on the average offsets per transient measured from the in vivo datasets. The range of the simulated spectral linewidth was determined from the median and upper bound, i.e., 95 percentile value, of the measured linewidth of the in vivo datasets. The range of the simulated spectral SNR was determined from the median and lower bound, i.e., 5 percentile value, of the measured spectral SNR of the in vivo datasets. These spectral parameters of the simulation represent the parameters of typical SVS studies at 7T. The values of the simulated spectral SNR (55.65-128) are within the ranges of the previously reported SNR of the SVS data at 7T (60.11-200) collected using different sequences and acquisition parameters [28,30]. The values of the simulated spectral linewidth (16-25 Hz) are broader than the previously reported linewidth (11.1 Hz) in the VOI of posterior cingulate cortex at 7T [28]. The broader simulated spectral linewidth in our study was to represent the linewidth measured in our selected VOI of anterior cingulate cortex, where the linewidth tends to be broader than the linewidth in other brain regions [31].

The conditions of the spectral linewidth and SNR considered in our study are within the ranges of typical SVS studies at 7T using different acquisition methods and VOIs on different brain regions. Different acquisition methods result in different impacts of macromolecule signals on the spectrum, which may influence the performance of the correction. The spectra collected with a short TE (TE<10 ms) [28] have a larger impact from macromolecule signals than the spectra collected with a TE of 30 ms in our study. The use of other water suppression methods [25], which generate stronger residual water signals as compared to the VAPOR method employed here, may benefit the RW method. Finally, the spectra in our study were unedited spectra and consisted of multiple metabolite peaks. For editing MR spectroscopy or multi-nuclear spectroscopy applications, the spectra could consist of only a few metabolite peaks, and the correction methods utilizing a single peak and multi-peak information could have similar performances.

The purpose of this study was to demonstrate the practical importance of each method by evaluating the changes in the spectral quality, similarity, and metabolite quantification after correction. The effect size estimates of the changes should provide informative assessment of the differences between the methods. Further statistical comparisons between methods were not performed in this study for two reasons. First, for practical importance, most of the methods showed a strong, moderate or weak effect of the correction-induced changes in our results. Further comparing two methods, which both showed a strong, moderate or weak effect, may not add to the practical importance of the methods. Second, our study was not designed for a fair comparison of the methods. We included the four methods which utilize different information of the spectrum and have different settings on the pre-processing procedure and frequency range to reduce effects from noise and unstable residual water signals. These settings have been shown to influence the performance of the methods [12,15,25]. However, in this study, we employed the previous settings at 3T without optimizing the settings for each method. For example, although the frequency ranges of the methods were largely outside the range of spurious echoes, we cannot eliminate the spurious echo effects which could be different among the methods. In view of these factors, we focused on reporting the effect size of the correction-induced changes for each method rather than rather than statistically comparing the changes between methods.

### Limitations

This study used the LCModel-simulated macromolecule spectra to model macromolecule signals, which only partially characterizes the macromolecule signals observed at 7T [45]. Given that the macromolecule pattern depends on B_0_, TE, TR, the acquisition sequence, subject’s age, and brain regions [45], previous studies have used the subject-specific macromolecule spectrum to account for the impact of macromolecules on the metabolite quantification [46–48]. However, it is challenging to perform subject-specific macromolecule measurements for large cohorts. An alternative approach is to parameterize the measured macromolecule spectrum into multiple components and include them in the basis set for the LCModel fit [47,48]. This approach requires a proper setting of parameterization and constraints to avoid overfitting [45]. Considering the above variables, the LCModel-simulated macromolecule spectra were employed for simplicity and reproducibility, although the incomplete characterization of macromolecules, e.g., in the range 3.4-4.1 ppm, may induce biases in the metabolite quantification. It remains to be investigated whether different approaches used to consider the macromolecule signals would influence the observed effects of the correction on the metabolite quantification. Another limitation is that the requirement of the 50 or more paired comparisons for the metabolite quantification analyses was an empirical selection, because the effect size was not known as prior information in this study.

## Conclusion

In the presence of the realistic frequency and phase offsets mainly induced by subject motion, the correction using each of the Cre, RW, SR, and Xcorr methods showed a strong to very strong effect of the improvements in the spectral linewidth, SNR and similarity. The correction only showed a weak to moderate effect of the changes in the metabolite quantification. Accordingly, each method provides practical benefits in improving spectral linewidth, SNR, and similarity. Whether these improvements translate into more reliable metabolite quantification at 7T requires further validation.

## A

**Supplementary Figure 1:** <u>Simulation:</u> (**a**) An example of applying the frequency and phase correction to the 32 simulated noise-free spectra of a dataset (spectral linewidth: 16 Hz) with added frequency and phase offsets. (**b**) The average errors of the frequency and phase measurements using each correction method were 0.05 Hz/0.5° (Cre), 0.05 Hz/0.01° (RW), 0.07 Hz/0.19° (SR), and 0.17 Hz/0.36° (Xcorr).

**Supplementary Figure 2:** A simulated noisy spectrum versus an in vivo spectrum (averaged over the 32 transients). The spectral linewidth and SNR were 16 Hz/128.29 of the simulated spectrum and were 17.64 Hz/125.39 of the in vivo spectrum.

**Supplementary Figure 3:** <u>Simulation:</u> The spectral linewidth, SNR, and SI_mean_ of the simulated spectrum (mean ± SD over the 20 repeated experiments) without and with the frequency and phase correction using each of the four methods under the specified conditions of the spectral SNR (**a**) and linewidth (**b**). The increased noise SD from 146 to 335 in **a** corresponds to a decrease in the spectral SNR from 128 to 55.65 with the line broadening factor (9.4 Hz) kept the same. The increased line broadening factor from 13.27 Hz to 22.27 Hz in **b** corresponds to an increase in the spectral linewidth from 16 to 25 Hz with the noise SD (146) kept the same.

**Supplementary Figure 4:** <u>Phantom data:</u> The measured frequency offset over 8 min and 32 s (mean over the 5 repeated experiments) on a phantom without prior MRI scans for 12 hours and following one-hour fMRI scan.

**Supplementary Figure 5:** <u>In vivo data:</u> Examples of the in vivo spectra (each with 32 transients) showing spurious echoes in the frequency ranges 4.1-4.5 ppm and 4.9-5.3 ppm.

**Supplementary Table 1:** <u>Simulation:</u> The concentrations (in mM) of the simulated signals used to synthesize the in vivo spectrum. The signals include 20 metabolites, residual water, and macromolecules. Their concentrations were determined from the median concentrations of the in vivo measurement from 127 participants. The zero concentration indicates that the metabolite signal cannot be detected due to a large relative uncertainty. MM refers to macromolecule resonances identified near 0.9 ppm (MM09), 2.0 ppm (MM20), 1.2 ppm (MM12), 1.4 ppm (MM14), and 1.7 ppm (MM17).

**Supplementary Table 2:** <u>Simulation:</u> The effect sizes and their 95% CIs of the spectral linewidth, SNR, and SI_mean_ changes following the frequency and phase correction using each of the four methods under the specified conditions of the spectral SNR as shown in Fig. 3a. The effect size was calculated using the Rosenthal formula. The 95% CI of the effect size was computed using bootstrap resampling with 10000 iterations.

**Supplementary Table 3:** <u>Simulation:</u> The effect sizes and their 95% CIs of the spectral linewidth, SNR, and SI_mean_ changes following the frequency and phase correction using each of the four methods under the specified conditions of the spectral linewidth as shown in Fig. 3b. The effect size was calculated using the Rosenthal formula. The 95% CI of the effect size was computed using bootstrap resampling with 10000 iterations.

## Notes

### Competing Interest Statement

V.A.M. receives research funding from GE Healthcare and SpinTech MRI.

### Funding

- National Institute of Health: R01MH111578, R01AG087159, S10RR028821

