## Supplementary for "Evaluations of retrospective frequency and phase correction methods for single-voxel MR spectroscopy at 7T"

### Slide 1
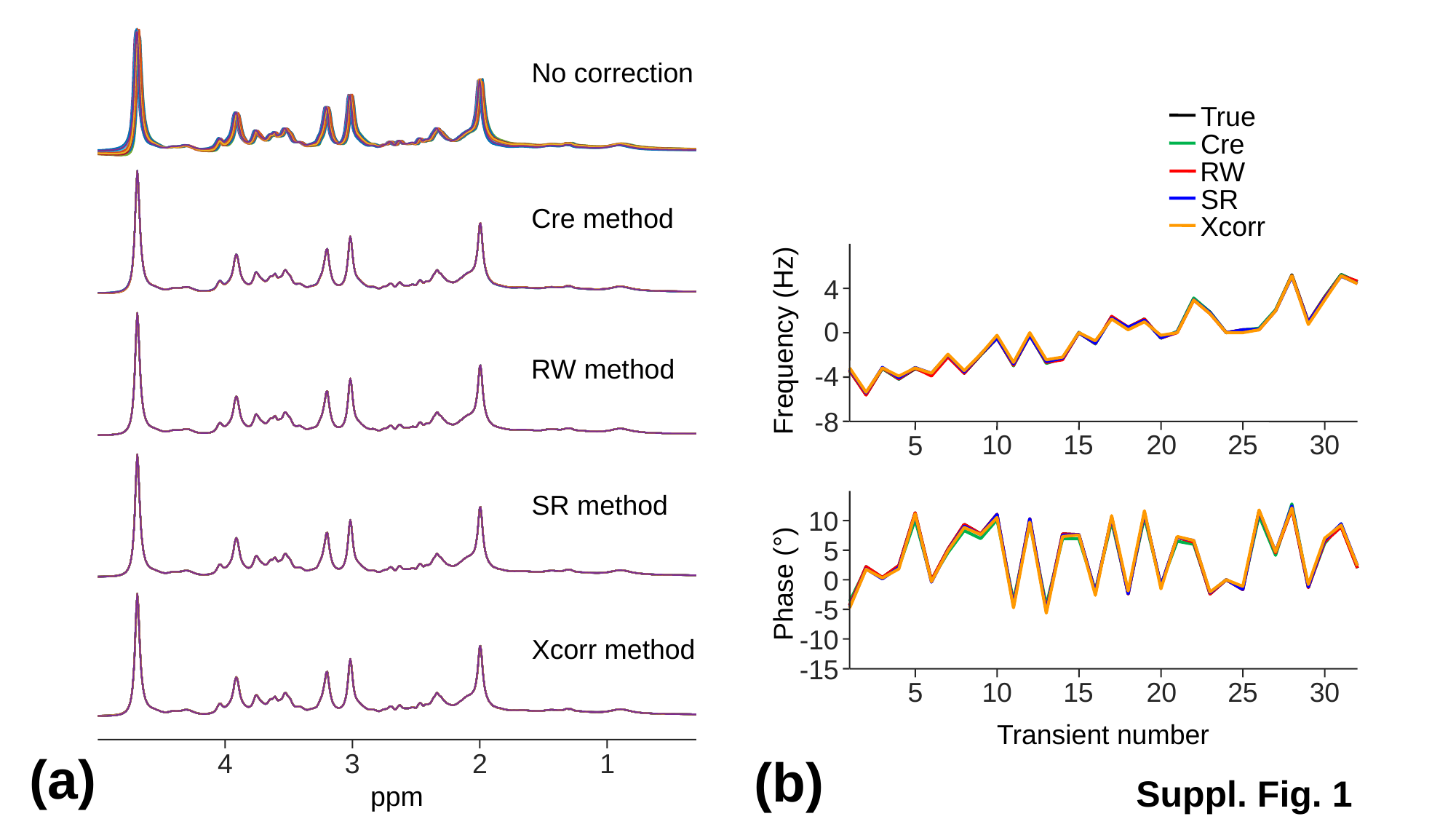

No correction
True
Cre
RW
SR
Xcorr
Cre method
4
0
Frequency (Hz)
-4
-8
RW method
25
30
20
10
15
5
SR method
10
5
0
Phase (°)
-5
-10
-15
Xcorr method
25
30
20
10
15
5
Transient number
(a)
4
3
2
1
(b)
Suppl. Fig. 1
ppm

### Slide 2
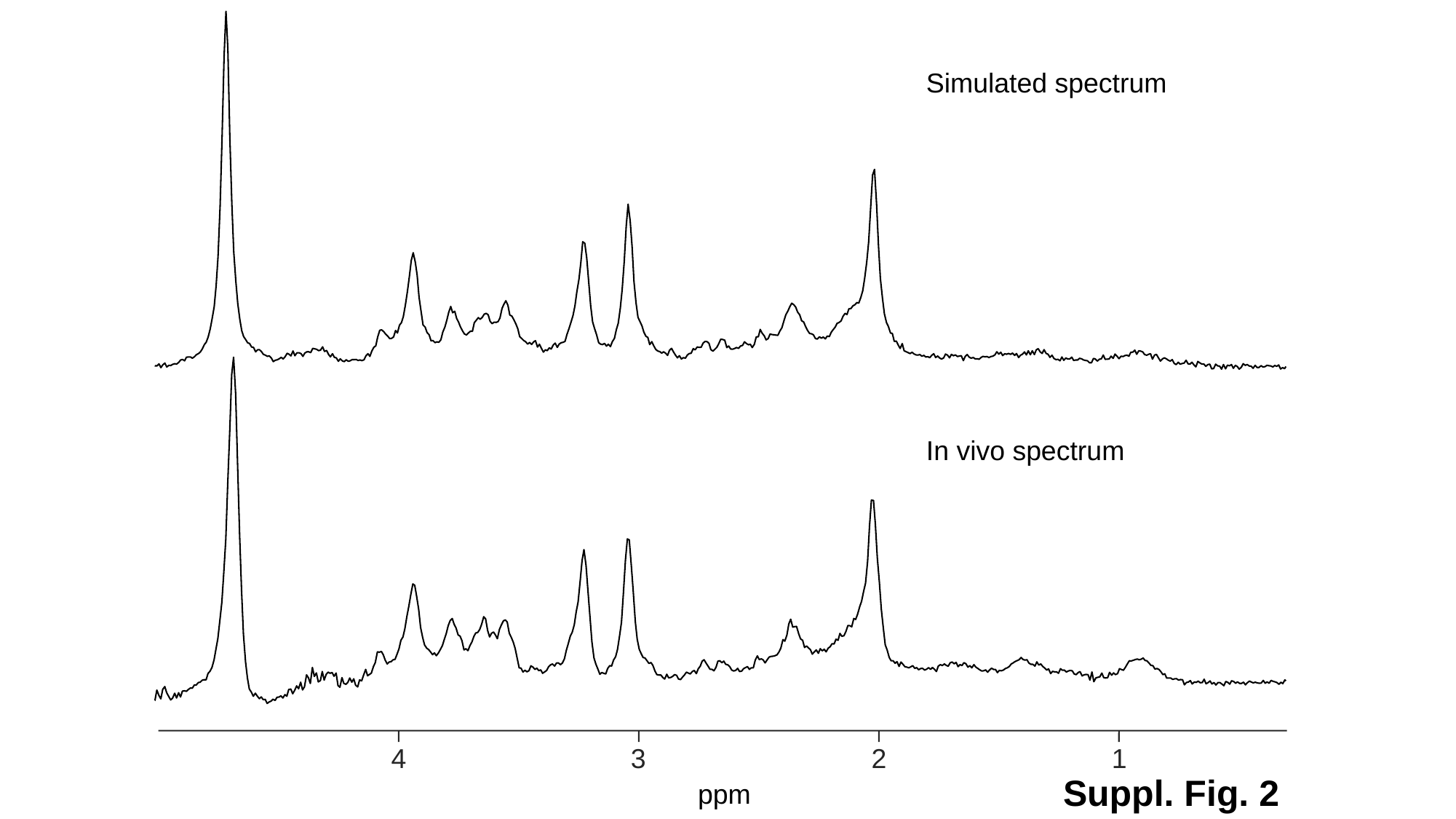

Simulated spectrum
In vivo spectrum
4
3
2
1
Suppl. Fig. 2
ppm

### Slide 3
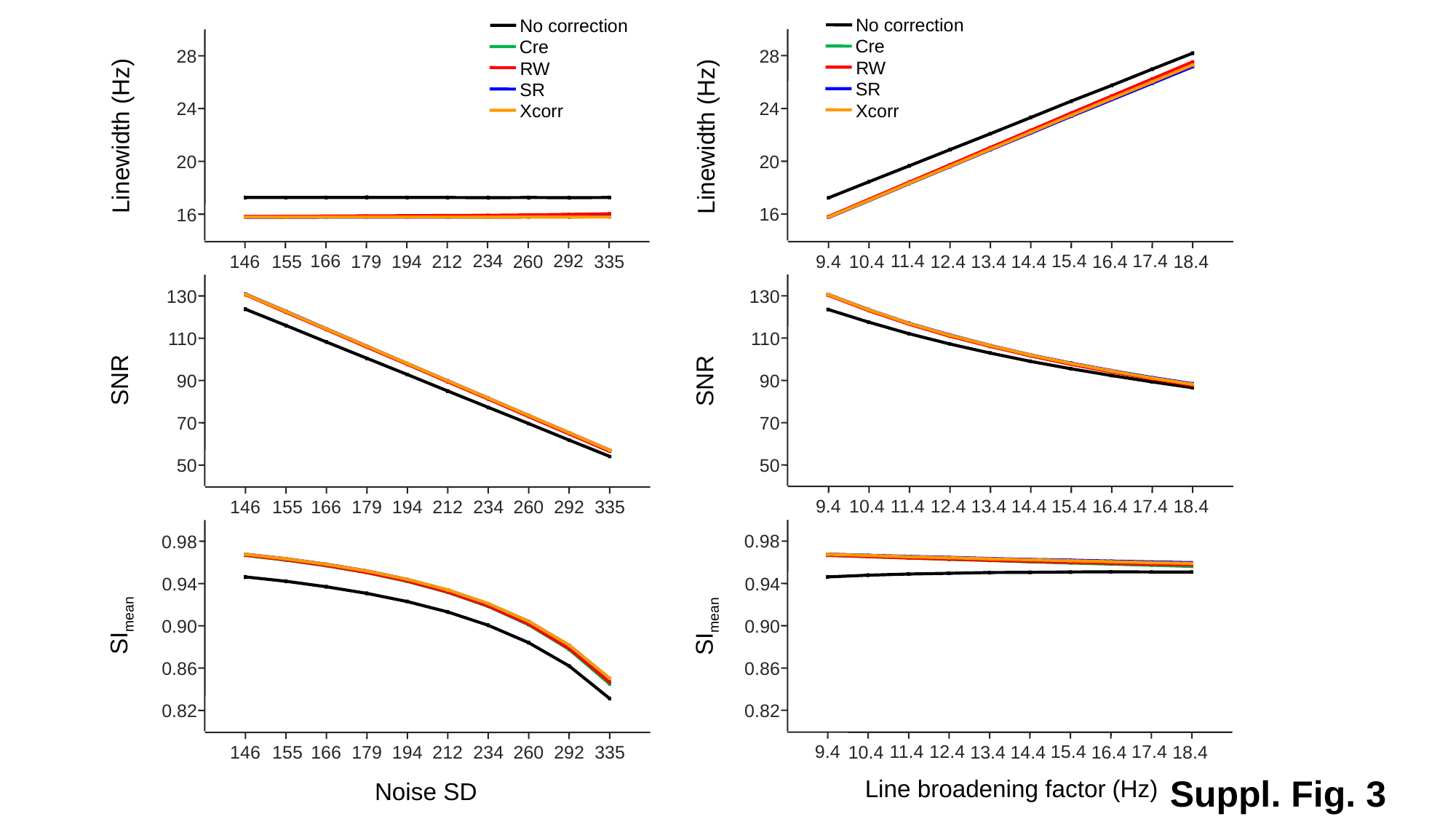

No correction
Cre
RW
SR
Xcorr
No correction
Cre
RW
SR
Xcorr
28
24
20
16
130
110
90
70
50
0.98
0.94
0.90
0.86
0.82
28
24
20
16
130
110
90
70
50
0.98
0.94
0.90
0.86
0.82
Linewidth (Hz)
Linewidth (Hz)
15.4
17.4
11.4
9.4
12.4
10.4
18.4
13.4
14.4
16.4
234
292
166
146
179
155
335
194
212
260
SNR
SNR
15.4
17.4
11.4
9.4
12.4
10.4
18.4
13.4
14.4
16.4
234
292
166
146
179
155
335
194
212
260
SI_mean_
SI_mean_
15.4
17.4
11.4
9.4
12.4
10.4
18.4
13.4
14.4
16.4
234
292
166
146
179
155
335
194
212
260
Suppl. Fig. 3
Line broadening factor (Hz)
Noise SD

### Slide 4
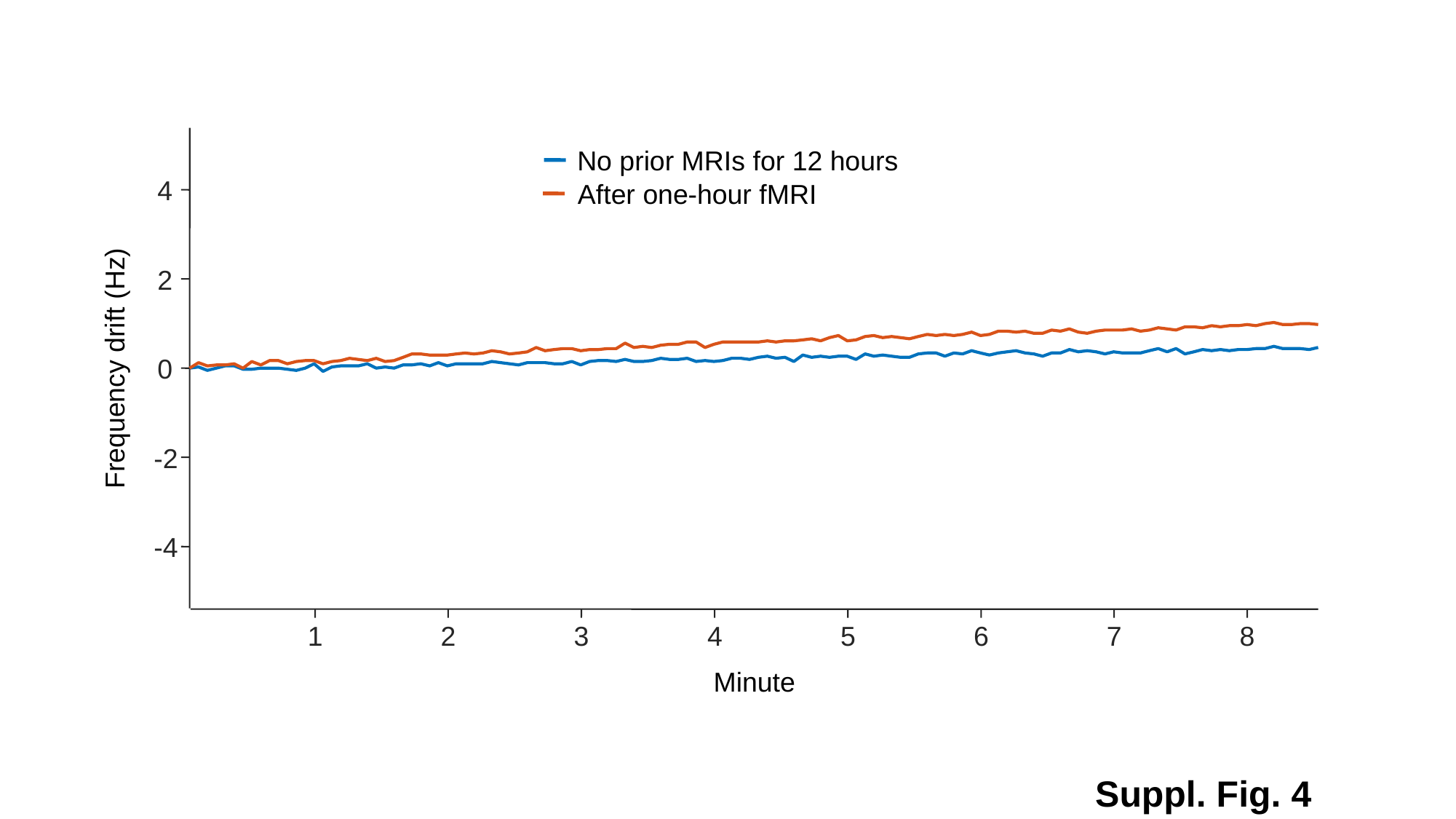

No prior MRIs for 12 hours
4
After one-hour fMRI
2
Frequency drift (Hz)
0
-2
-4
7
8
1
2
3
4
5
6
Minute
Suppl. Fig. 4

### Slide 5
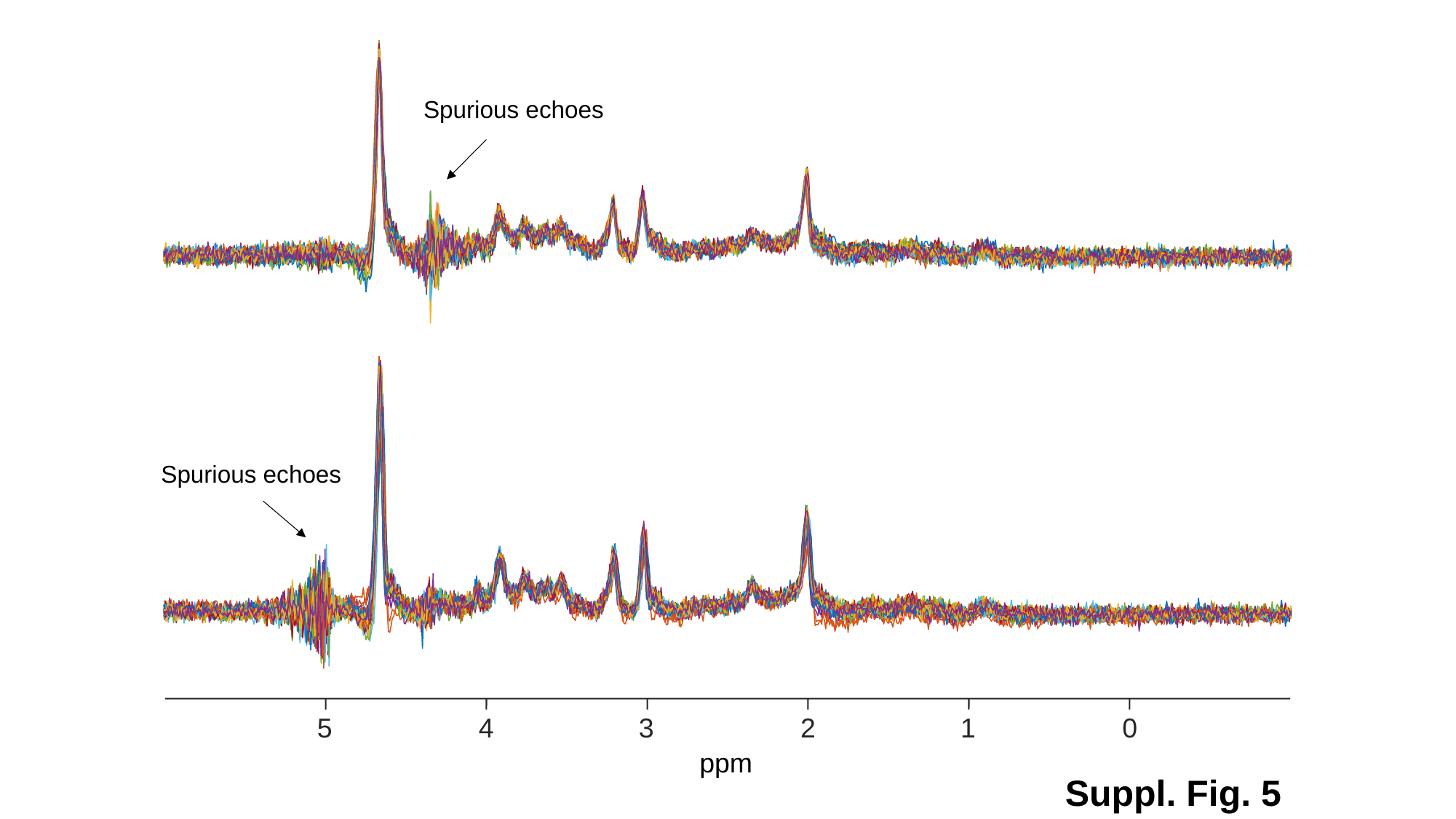

Spurious echoes
Spurious echoes
5
4
3
2
1
0
ppm
Suppl. Fig. 5

### Slide 6
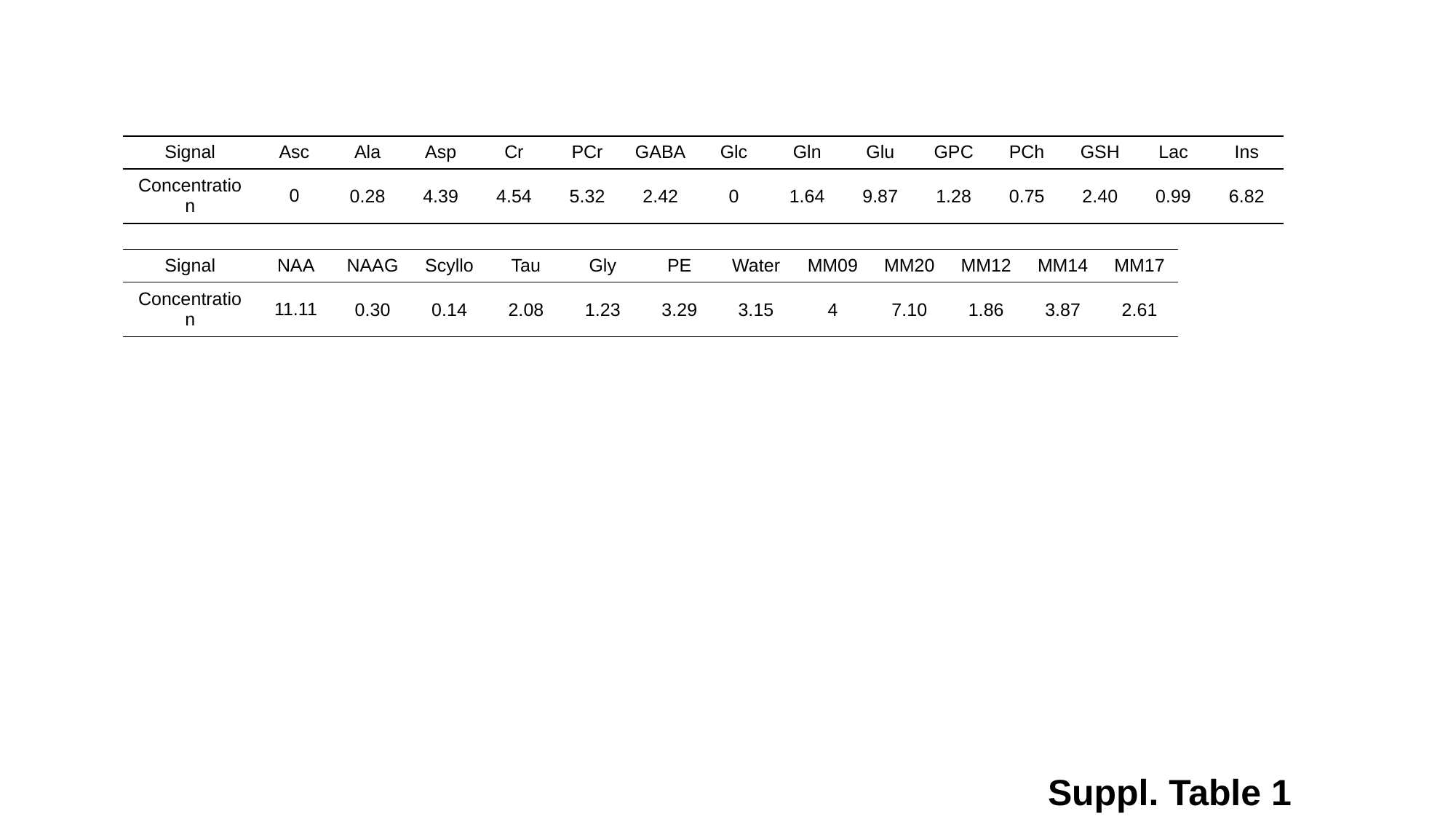

| Signal | Asc | Ala | Asp | Cr | PCr | GABA | Glc | Gln | Glu | GPC | PCh | GSH | Lac | Ins |
| --- | --- | --- | --- | --- | --- | --- | --- | --- | --- | --- | --- | --- | --- | --- |
| Concentration | 0 | 0.28 | 4.39 | 4.54 | 5.32 | 2.42 | 0 | 1.64 | 9.87 | 1.28 | 0.75 | 2.40 | 0.99 | 6.82 |
| Signal | NAA | NAAG | Scyllo | Tau | Gly | PE | Water | MM09 | MM20 | MM12 | MM14 | MM17 |
| --- | --- | --- | --- | --- | --- | --- | --- | --- | --- | --- | --- | --- |
| Concentration | 11.11 | 0.30 | 0.14 | 2.08 | 1.23 | 3.29 | 3.15 | 4 | 7.10 | 1.86 | 3.87 | 2.61 |
Suppl. Table 1

### Slide 7
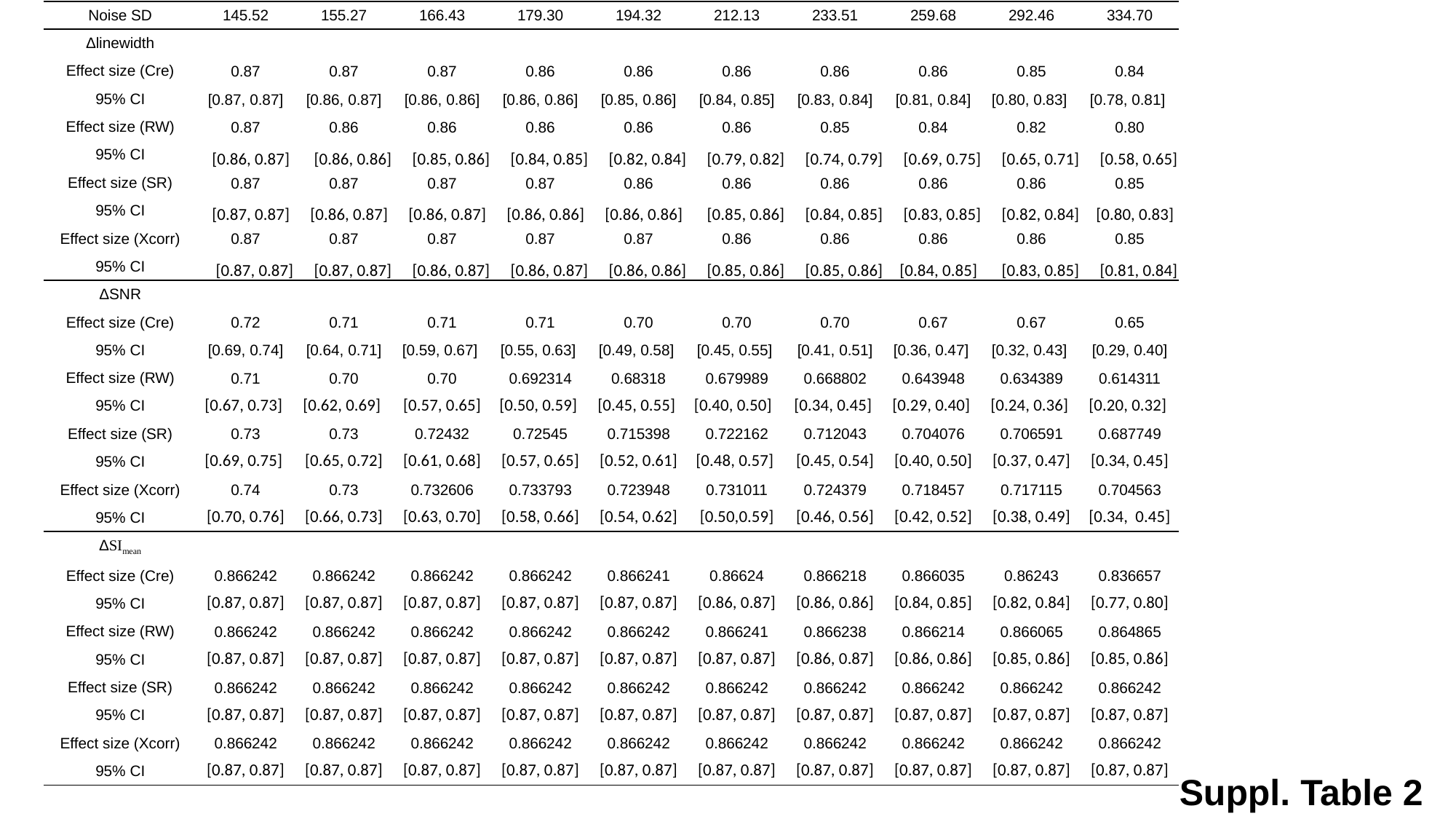

| Noise SD | 145.52 | 155.27 | 166.43 | 179.30 | 194.32 | 212.13 | 233.51 | 259.68 | 292.46 | 334.70 |
| --- | --- | --- | --- | --- | --- | --- | --- | --- | --- | --- |
| Δlinewidth | | | | | | | | | | |
| Effect size (Cre) | 0.87 | 0.87 | 0.87 | 0.86 | 0.86 | 0.86 | 0.86 | 0.86 | 0.85 | 0.84 |
| 95% CI | [0.87, 0.87] | [0.86, 0.87] | [0.86, 0.86] | [0.86, 0.86] | [0.85, 0.86] | [0.84, 0.85] | [0.83, 0.84] | [0.81, 0.84] | [0.80, 0.83] | [0.78, 0.81] |
| Effect size (RW) | 0.87 | 0.86 | 0.86 | 0.86 | 0.86 | 0.86 | 0.85 | 0.84 | 0.82 | 0.80 |
| 95% CI | [0.86, 0.87] | [0.86, 0.86] | [0.85, 0.86] | [0.84, 0.85] | [0.82, 0.84] | [0.79, 0.82] | [0.74, 0.79] | [0.69, 0.75] | [0.65, 0.71] | [0.58, 0.65] |
| Effect size (SR) | 0.87 | 0.87 | 0.87 | 0.87 | 0.86 | 0.86 | 0.86 | 0.86 | 0.86 | 0.85 |
| 95% CI | [0.87, 0.87] | [0.86, 0.87] | [0.86, 0.87] | [0.86, 0.86] | [0.86, 0.86] | [0.85, 0.86] | [0.84, 0.85] | [0.83, 0.85] | [0.82, 0.84] | [0.80, 0.83] |
| Effect size (Xcorr) | 0.87 | 0.87 | 0.87 | 0.87 | 0.87 | 0.86 | 0.86 | 0.86 | 0.86 | 0.85 |
| 95% CI | [0.87, 0.87] | [0.87, 0.87] | [0.86, 0.87] | [0.86, 0.87] | [0.86, 0.86] | [0.85, 0.86] | [0.85, 0.86] | [0.84, 0.85] | [0.83, 0.85] | [0.81, 0.84] |
| ΔSNR | | | | | | | | | | |
| Effect size (Cre) | 0.72 | 0.71 | 0.71 | 0.71 | 0.70 | 0.70 | 0.70 | 0.67 | 0.67 | 0.65 |
| 95% CI | [0.69, 0.74] | [0.64, 0.71] | [0.59, 0.67] | [0.55, 0.63] | [0.49, 0.58] | [0.45, 0.55] | [0.41, 0.51] | [0.36, 0.47] | [0.32, 0.43] | [0.29, 0.40] |
| Effect size (RW) | 0.71 | 0.70 | 0.70 | 0.692314 | 0.68318 | 0.679989 | 0.668802 | 0.643948 | 0.634389 | 0.614311 |
| 95% CI | [0.67, 0.73] | [0.62, 0.69] | [0.57, 0.65] | [0.50, 0.59] | [0.45, 0.55] | [0.40, 0.50] | [0.34, 0.45] | [0.29, 0.40] | [0.24, 0.36] | [0.20, 0.32] |
| Effect size (SR) | 0.73 | 0.73 | 0.72432 | 0.72545 | 0.715398 | 0.722162 | 0.712043 | 0.704076 | 0.706591 | 0.687749 |
| 95% CI | [0.69, 0.75] | [0.65, 0.72] | [0.61, 0.68] | [0.57, 0.65] | [0.52, 0.61] | [0.48, 0.57] | [0.45, 0.54] | [0.40, 0.50] | [0.37, 0.47] | [0.34, 0.45] |
| Effect size (Xcorr) | 0.74 | 0.73 | 0.732606 | 0.733793 | 0.723948 | 0.731011 | 0.724379 | 0.718457 | 0.717115 | 0.704563 |
| 95% CI | [0.70, 0.76] | [0.66, 0.73] | [0.63, 0.70] | [0.58, 0.66] | [0.54, 0.62] | [0.50,0.59] | [0.46, 0.56] | [0.42, 0.52] | [0.38, 0.49] | [0.34, 0.45] |
| ΔSI_mean_ | | | | | | | | | | |
| Effect size (Cre) | 0.866242 | 0.866242 | 0.866242 | 0.866242 | 0.866241 | 0.86624 | 0.866218 | 0.866035 | 0.86243 | 0.836657 |
| 95% CI | [0.87, 0.87] | [0.87, 0.87] | [0.87, 0.87] | [0.87, 0.87] | [0.87, 0.87] | [0.86, 0.87] | [0.86, 0.86] | [0.84, 0.85] | [0.82, 0.84] | [0.77, 0.80] |
| Effect size (RW) | 0.866242 | 0.866242 | 0.866242 | 0.866242 | 0.866242 | 0.866241 | 0.866238 | 0.866214 | 0.866065 | 0.864865 |
| 95% CI | [0.87, 0.87] | [0.87, 0.87] | [0.87, 0.87] | [0.87, 0.87] | [0.87, 0.87] | [0.87, 0.87] | [0.86, 0.87] | [0.86, 0.86] | [0.85, 0.86] | [0.85, 0.86] |
| Effect size (SR) | 0.866242 | 0.866242 | 0.866242 | 0.866242 | 0.866242 | 0.866242 | 0.866242 | 0.866242 | 0.866242 | 0.866242 |
| 95% CI | [0.87, 0.87] | [0.87, 0.87] | [0.87, 0.87] | [0.87, 0.87] | [0.87, 0.87] | [0.87, 0.87] | [0.87, 0.87] | [0.87, 0.87] | [0.87, 0.87] | [0.87, 0.87] |
| Effect size (Xcorr) | 0.866242 | 0.866242 | 0.866242 | 0.866242 | 0.866242 | 0.866242 | 0.866242 | 0.866242 | 0.866242 | 0.866242 |
| 95% CI | [0.87, 0.87] | [0.87, 0.87] | [0.87, 0.87] | [0.87, 0.87] | [0.87, 0.87] | [0.87, 0.87] | [0.87, 0.87] | [0.87, 0.87] | [0.87, 0.87] | [0.87, 0.87] |
Suppl. Table 2

### Slide 8
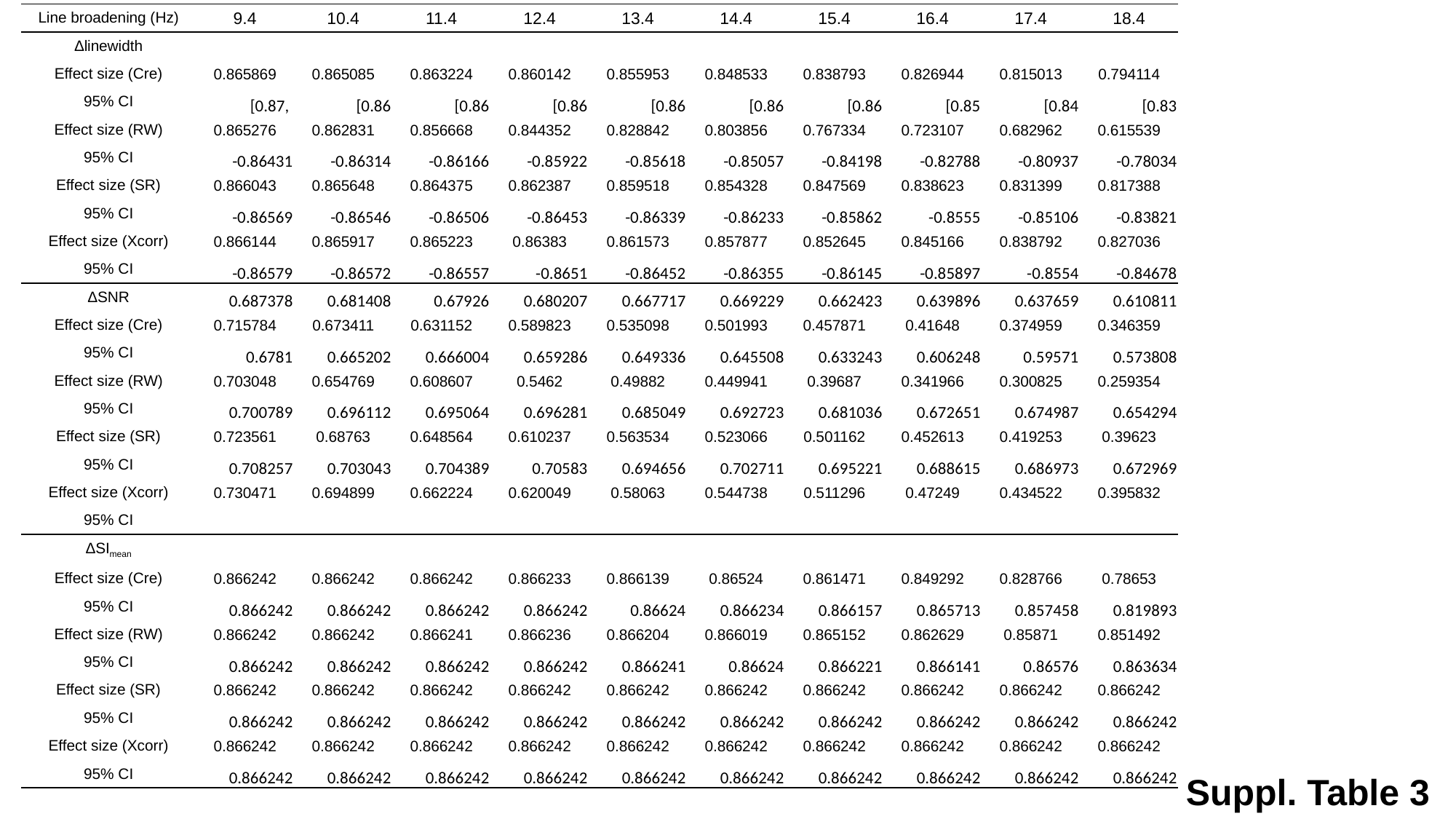

| Line broadening (Hz) | 9.4 | 10.4 | 11.4 | 12.4 | 13.4 | 14.4 | 15.4 | 16.4 | 17.4 | 18.4 |
| --- | --- | --- | --- | --- | --- | --- | --- | --- | --- | --- |
| Δlinewidth | | | | | | | | | | |
| Effect size (Cre) | 0.865869 | 0.865085 | 0.863224 | 0.860142 | 0.855953 | 0.848533 | 0.838793 | 0.826944 | 0.815013 | 0.794114 |
| 95% CI | [0.87, | [0.86 | [0.86 | [0.86 | [0.86 | [0.86 | [0.86 | [0.85 | [0.84 | [0.83 |
| Effect size (RW) | 0.865276 | 0.862831 | 0.856668 | 0.844352 | 0.828842 | 0.803856 | 0.767334 | 0.723107 | 0.682962 | 0.615539 |
| 95% CI | -0.86431 | -0.86314 | -0.86166 | -0.85922 | -0.85618 | -0.85057 | -0.84198 | -0.82788 | -0.80937 | -0.78034 |
| Effect size (SR) | 0.866043 | 0.865648 | 0.864375 | 0.862387 | 0.859518 | 0.854328 | 0.847569 | 0.838623 | 0.831399 | 0.817388 |
| 95% CI | -0.86569 | -0.86546 | -0.86506 | -0.86453 | -0.86339 | -0.86233 | -0.85862 | -0.8555 | -0.85106 | -0.83821 |
| Effect size (Xcorr) | 0.866144 | 0.865917 | 0.865223 | 0.86383 | 0.861573 | 0.857877 | 0.852645 | 0.845166 | 0.838792 | 0.827036 |
| 95% CI | -0.86579 | -0.86572 | -0.86557 | -0.8651 | -0.86452 | -0.86355 | -0.86145 | -0.85897 | -0.8554 | -0.84678 |
| ΔSNR | 0.687378 | 0.681408 | 0.67926 | 0.680207 | 0.667717 | 0.669229 | 0.662423 | 0.639896 | 0.637659 | 0.610811 |
| Effect size (Cre) | 0.715784 | 0.673411 | 0.631152 | 0.589823 | 0.535098 | 0.501993 | 0.457871 | 0.41648 | 0.374959 | 0.346359 |
| 95% CI | 0.6781 | 0.665202 | 0.666004 | 0.659286 | 0.649336 | 0.645508 | 0.633243 | 0.606248 | 0.59571 | 0.573808 |
| Effect size (RW) | 0.703048 | 0.654769 | 0.608607 | 0.5462 | 0.49882 | 0.449941 | 0.39687 | 0.341966 | 0.300825 | 0.259354 |
| 95% CI | 0.700789 | 0.696112 | 0.695064 | 0.696281 | 0.685049 | 0.692723 | 0.681036 | 0.672651 | 0.674987 | 0.654294 |
| Effect size (SR) | 0.723561 | 0.68763 | 0.648564 | 0.610237 | 0.563534 | 0.523066 | 0.501162 | 0.452613 | 0.419253 | 0.39623 |
| 95% CI | 0.708257 | 0.703043 | 0.704389 | 0.70583 | 0.694656 | 0.702711 | 0.695221 | 0.688615 | 0.686973 | 0.672969 |
| Effect size (Xcorr) | 0.730471 | 0.694899 | 0.662224 | 0.620049 | 0.58063 | 0.544738 | 0.511296 | 0.47249 | 0.434522 | 0.395832 |
| 95% CI | | | | | | | | | | |
| ΔSI_mean_ | | | | | | | | | | |
| Effect size (Cre) | 0.866242 | 0.866242 | 0.866242 | 0.866233 | 0.866139 | 0.86524 | 0.861471 | 0.849292 | 0.828766 | 0.78653 |
| 95% CI | 0.866242 | 0.866242 | 0.866242 | 0.866242 | 0.86624 | 0.866234 | 0.866157 | 0.865713 | 0.857458 | 0.819893 |
| Effect size (RW) | 0.866242 | 0.866242 | 0.866241 | 0.866236 | 0.866204 | 0.866019 | 0.865152 | 0.862629 | 0.85871 | 0.851492 |
| 95% CI | 0.866242 | 0.866242 | 0.866242 | 0.866242 | 0.866241 | 0.86624 | 0.866221 | 0.866141 | 0.86576 | 0.863634 |
| Effect size (SR) | 0.866242 | 0.866242 | 0.866242 | 0.866242 | 0.866242 | 0.866242 | 0.866242 | 0.866242 | 0.866242 | 0.866242 |
| 95% CI | 0.866242 | 0.866242 | 0.866242 | 0.866242 | 0.866242 | 0.866242 | 0.866242 | 0.866242 | 0.866242 | 0.866242 |
| Effect size (Xcorr) | 0.866242 | 0.866242 | 0.866242 | 0.866242 | 0.866242 | 0.866242 | 0.866242 | 0.866242 | 0.866242 | 0.866242 |
| 95% CI | 0.866242 | 0.866242 | 0.866242 | 0.866242 | 0.866242 | 0.866242 | 0.866242 | 0.866242 | 0.866242 | 0.866242 |
Suppl. Table 3
